# Capsid Proteins are Necessary for Replication of a Parvovirus

**DOI:** 10.1101/2020.04.06.028852

**Authors:** Thomas Labadie, Deborah Garcia, Doriane Mutuel, Mylène Ogliastro, Guillaume Cambray

**Affiliations:** Diversité des Génomes et Interactions Microorganismes Insectes (DGIMI), UMR 1333, Univ Montpellier, INRA, 34095 Montpellier, France; Centre de Biochimie Structurale (CBS), UMR 5048, Univ Montpellier, CNRS, 34090 Montpellier, France

## Abstract

Despite tight genetic compression, viral genomes are often organized in functional gene clusters, a modular structure that might favor their evolvability. This has greatly facilitated biotechnological developments, such as the recombinant Adeno-Associated Virus (AAV) systems for gene therapy. Following this lead, we endeavored to engineer the related insect parvovirus *Junonia coenia* densovirus (JcDV) to create addressable vectors for insect pest biocontrol. To enable safer manipulation of capsid mutants, we translocated the non-structural (*ns*) gene cluster outside the viral genome. To our dismay, this yielded a virtually non-replicable clone. We linked the replication defect to an unexpected modularity breach, as *ns* translocation truncated the overlapping 3’ UTR of the capsid transcript (*vp*). We found that native *vp* 3’UTR is necessary to high VP production, but that decreased expression do not adversely impact the expression of NS proteins, which are known replication effectors. As nonsense *vp* mutations recapitulate the replication defect, VP proteins appear directly implicated in the replication process. Our findings suggest intricate replication-encapsidation couplings that favor maintenance of genetic integrity. We discuss possible connections with an intriguing cis-packaging phenomenon previously observed in parvoviruses, whereby capsids preferentially package the genome from which they were expressed.

**Importance:** Densoviruses could be used as biological control agents to manage insect pests. Such applications require in depth biological understanding and associated molecular tools. However, the genomes of these viruses remain hard to manipulate due too poorly tractable secondary structures at their extremities. We devised a construction strategy that enable precise and efficient molecular modifications. Using this approach, we endeavored to create a split clone of the *Junonia coenia* densovirus (JcDV) that can be used to safely study the impact of capsid mutations on host specificity. Our original construct proved to be non-functional. Fixing this defect led us to uncover that capsid proteins and their correct expression are essential for continued rolling-hairpin replication. This points to an intriguing link between replication and packaging, which might be shared with related viruses. This serendipitous discovery illustrates the power of synthetic biology approaches to advance our knowledge of biological systems.

## Introduction

Viruses represent the most diverse and abundant biological entities on earth and can be found in nearly all organisms. Over the last decade, it has become increasingly apparent that the evolutionary success of viruses is largely supported by their propensity to recombine (1). As if to promote evolvability, the genome of many viruses displays modular architecture, wherein sequences involved with capsid formation and those coding for replicative functions are segregated in distinct functional blocks.

Such modularity is clearly apparent in *Parvoviridae*. Member of this family are characterized by single stranded DNA genomes of 4-6kb that invariably consist in similarly sized *vp* and *ns* gene clusters, flanked by telomeric sequences. The *vp* (viral particle) block bears 1-4 sequences coding for an icosahedral capsid (T=1), while the *ns* (non-structural) block code for 1-3 proteins involved in replication, packaging and other often undefined functions. Telomeric sequences can fold into hairpins involved in replication and also contain packaging signals. This modular organization has been exploited in the context of gene therapy to, for example, produce pseudotyped recombinant AAV virions based on telomeres, *ns* and *vp* genes from different parvoviruses (2).

In this work, we wanted to physically dissociate the *vp* and *ns* gene clusters to generate a non-propagatable clone of the Junonia Coenia Densovirus (JcDV).

JcDV is the type member of the genus *Protoambidensovirus*, which comprises invertebrate parvoviruses characterized by tail to tail orientation of their *ns* and *vp* blocks (Figure 1A). This original architecture usually results in a 60 nts overlap between the 3’UTR of their respective transcripts (3). JcDV can be pathogenic for several species of insects from the *Lepidoptera* order, including agriculturally relevant pests such as the Fall Army Worm (*Spodoptera frugiperda*), a polyphagous crop pest that is currently spreading over the world at alarming rates (4). Densoviruses have long been considered promising candidates for the biological control of insect populations, and several field trials have confirmed their potential efficacies (5, 6).

**Figure 1.**
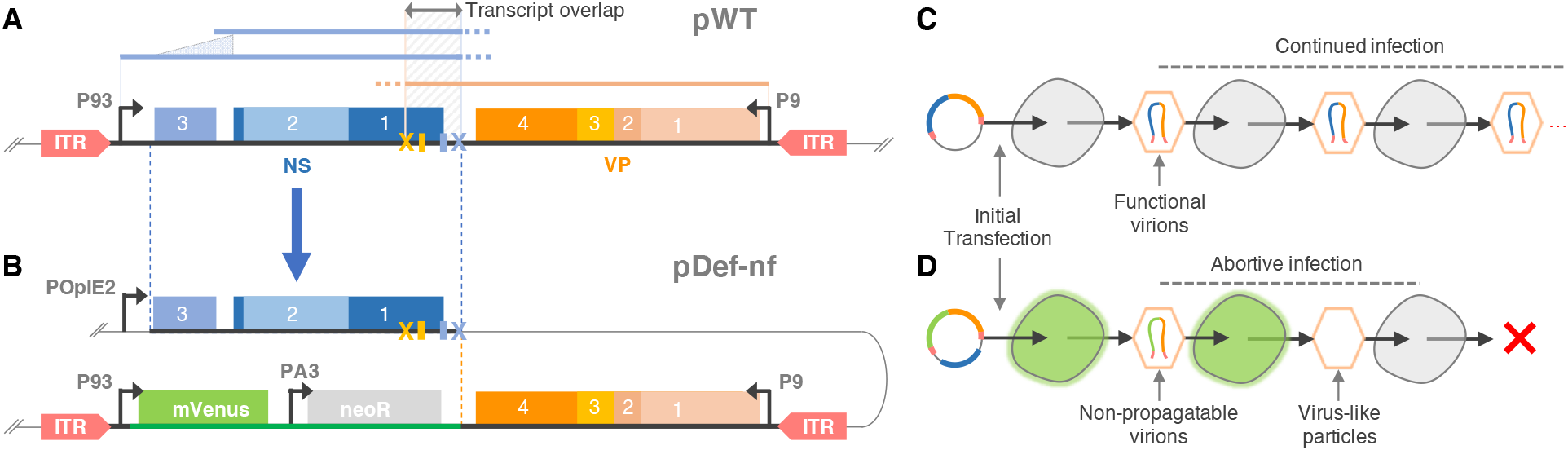
Initial design for the construction of a non-propagatable JcDV clone. **A.** Functional architecture of the infectious JcDV clone pWT. *ns* (blue) and *vp* (orange) gene blocks are in tail to tail orientation. Each produce a single transcript (blue and orange lines) from a promoter partly located in the inverted terminal repeats (ITRs, red). The *ns* transcript is alternatively spliced, as shown. Due to the location of *ns* and *vp* polyadenylation signals and sites (blue and orange small rectangle and cross, respectively), the two transcripts overlap over a complementary region of 60 nucleotides (grey arrow, top). **B.** Architecture of the non-propagatable clone pDef-nf. The *ns* block (start of *ns3* to end of *ns* transcript) was cloned out of the genome under the control of a strong baculovirus promoter, and replaced by a reporter block of the same size comprising the *mVenus* fluorescent protein gene and a neomycin resistance gene. In this process, VP’s 3’UTR was truncated by 44 nts to avoid recombination-prone sequence duplication. **C.** Continued infection upon transfection of the pWT clone. Upon packaging of the WT genome, produced virions can propagate to naïve cells leading to uncontrolled spread of the infection. **D.** Transfection of the non-propagatable pDef-nf clone should not permit downstream infection. *Trans* complementation by NS function enable initial genome replication in transfected cells. Upon packaging, genomes devoid of NS functions cannot replicate in naïve cells, containing the spread of the virus.

In this perspective, the ability to fine tune host tropism would be particularly desirable to establish precise and controllable biocontrol solutions. As with other vertebrate parvoviruses (7, 8), the variation of few amino-acid residues at the surface of JcDV’s capsid have been shown to modulate JcDV’s host range (9). To better understand—and eventually control—these sequence determinants, we endeavored to develop a high-throughput screen of rationally engineered capsid variants in which the link between mutations and consequent capsid performances is established by deep sequencing of mutated *vp* genes packaged in their cognate capsid (10–12). Concerned with potential biohazards, we first wished to establish a non-propagatable system wherein mutant virions could be screened for capsid functionality, while not being capable of effecting further productive infection and transmission.

To produce non-propagatable virions that can nonetheless package the *vp* gene required for subsequent sequencing-based capsid identification, we moved the *ns* gene block in *trans* and replaced it by a reporter gene block of the same size. Although similar manipulation had been successfully accomplished for several parvoviruses, ours yielded a completely nonfunctional JcDV clone. Subsequent investigation led us to link the alteration of *vp* transcript’s 3’UTR to a stark decrease in VP protein production and a near complete shutdown of viral replication. This allowed us to unveil a direct and essential involvement of the VP proteins in the replication process of a parvovirus.

## Material and Methods

### Plasmid constructions

All enzymes were purchased from NEB. Phusion Hot-Start Flex polymerase (NEB) was used for all PCR reactions. NEB Turbo cells were used for transformation of all constructs, except construct comprising ITRs, which were transformed in NEB stable strain and grown at 30°C to improve genetic stability. Primers were ordered from IDT. Constructions were systematically verified by Sanger sequencing (Eurogentec). Constructions assayed in this work are listed in Table 1. A more exhaustive plasmid list including construction intermediates described below is provided in Supplementary Table 1. Oligonucleotide sequences are listed in Supplementary Table 2.

**Table 1.** Plasmids used in this work.

| Name | Description | ID | Reference |
| --- | --- | --- | --- |
| pWT | WT JcDV clone derived from the original pBRJH clone | p14 | This work |
| pDef-nF | Original (non-functional) design of JcDV's non-propagatable clone with 16 nts after *vp*'s stop codon | p58 | This work |
| pNS | JcDV's P93 and *ns* followed by 484 nts after *ns1*'s stop codon | p90 | 17 |
| pVP | JcDV's P9 and *vp* followed by 115 nts after *vp*'s stop codon | p91 | 17 |
| pWT-ITRless | pBRJH derivative with truncated ITRs (no hairpin, no replication) | p92 | 16 |
| pNS-3p49 | JcDV's P9 and *vp* followed by 49 nts after *vp*'s stop codon | p93 | This work |
| pDef-3p115 | Amended (functional) design of JcDV's non-propagatable clone with 115 nts after *vp*'s stop codon | p103 | This work |
| pDef-3p91 | Amended (functional) design of JcDV's non-propagatable clone with 91 nts after *vp*'s stop codon | p104 | This work |
| pITR-R | JcDV's extended (oxford) ITRs flanking a DNA sequence of the same length as the WT genome. Comprises *eGFP* driven by *B. Mori*'s A3 promoter and followed by SV40 polyadenylation signal | p116 | This work |
| pVP-3p75 | JcDV's P9 and *vp* followed by 75 nts after *vp*'s stop codon (stops just after polyA site) | p118 | This work |
| pVP-7m | JcDV's P9 and *vp* followed by 115 nts after *vp*'s stop codon with 7 substitutions in the *ns1* overlapping region | p144 | This work |
| pVP-12m | JcDV's P9 and *vp* followed by 115 nts after *vp*'s stop codon with 12 substitutions in the *ns1* overlapping region | p145 | This work |
| pNS-7m | JcDV's P93 and *ns* followed by 484 nts after *ns1*'s stop codon. 7 silent substitutions in the end of *ns1* | p146 | This work |
| pNS-12m | JcDV's P93 and *ns* followed by 484 nts after *ns1*'s stop codon. 12 silent substitutions in the end of *ns1* | p147 | This work |
| pDef-3p115-S | Amended design of JcDV's non-propagatable clone with 115 nts after *vp*'s stop codon and TAT 542 TAG substitution in *vp* (amber stop codon) | p254 | This work |
| pDef-3p115-F | Amended design of JcDV's non-propagatable clone with 115 nts after *vp*'s stop codon and AAT 687 AA- deletion in *vp* (frameshift) | p255 | This work |

#### Construction of JcDV clones

The WT JcDV clone (pWT) was derived from the published pBRJH clone (13) by outcloning the *cat* cassette (conferring resistance to chloramphenicol) from the pSB1C3 of p10 in place of the *bla* cassette (conferring resistance to ampicillin) upon BspHI digest. Non-propagatable vectors were constructed using Golden Gate cloning (14) of several fragments to avoid problematic PCR amplification of unstable ITRs. A graphical overview is presented in Supplementary Figure 1. Fragments were flanked by BsaI sites to eventually enable directed and seamless cloning and initially subcloned into a minimal donor backbone comprising a p15a origin of replication and chloramphenicol resistance (pBbA2c-RFP(15) modified to contain matching BsaI sites. An extended version of JcDV’s ITR (16) was synthesized (Genewiz, p39) and subcloned into the donor vector amplified by o122+o147 to yield p47. The P93 promoter was amplified from pWT (o114+115) and cloned into the donor vector amplified by o127+o128 to yield p46. A reporter block comprised of eGFP, actin promoter A3 from *Bombyx mori* and a neomycin resistance gene (*neo*) was constructed by assembly PCR. eGFP and PA3 were separately amplified from pITR-A3-GFP (13) using o54+o6 and o7+o8, respectively. The *neo* gene was amplified from pNEO (Amersham) using o9+o55. The 3 amplicons were assembled using o54+o55 and cloned into the donor vector amplified by o56+o57 to yield p17. eGFP was then substituted by mVenus: p17 was amplified with o63+o64; mVenus was amplified from plasmid mCerulean3-mVenus-FRET-10 (a gift from Michael Davidson, Addgene plasmid # 58180) using o61+o62; and the two amplicons were cloned at BsmBI sites introduced during the PCRs to yield p36. The sequence corresponding to P9-vp123 (*i.e.* excluding the *vp4* part) was amplified from pWT using o114+o94 and cloned into the donor backbone amplified with o85+127 to yield p42. In this process, a double BsmBI site was introduced to enable subsequent in-frame cloning of *vp4*. The latter was amplified from pWT using o96+o97 and cloned into the donor backbone amplified with o116+o117 to yield p40. Unlike other donor constructs, *vp4* in p40 is flanked by BsmBI sites. To construct the acceptor plasmid, we first cloned the *ns* gene block under the control of a strong OpEI-2 baculoviral promoter in a pIZ backbone. For this, we amplified the *ns* gene block using an assembly PCR designed to remove a BsaI site in *ns3* through the introduction of a silent Arg substitution (AG<u>G</u>119AG<u>A</u>). The two *ns* fragments were amplified from pWT using o112+o19 and o113+o20, assembled using o112+113 and cloned between BamHI and AgeI sites of p25 (pIZ-Flag6His-Siwi, (17) to yield p45. A landing pad comprised of two BsaI sites was assembled by annealing o120+o121 and cloned at the BspHI site of p45 to yield p49.

Donor vectors p47 (ITR), p46 (P93), p36 (mVenus-A3-neo) and p42 (P9-vp123-2xBsmBI) and acceptor vector p49 were used in a one-pot Golden Gate reaction with BsaI to yield the *vp4*-less intermediate p57. The non-propagatable clone pDef-nf was constructed by Golden Gate cloning of p40 into p57 using BsmBI.

To construct amended versions of this non-functional clone, longer versions of p40 that extend the sequence 3’ of vp4 were amplified from pWT using o243+o239 and o243+244 and cloned into the donor backbone amplified by o116+o117 to yield p105 and p106, respectively. These were introduced in p57 by Golden Gate cloning (BsmBI) to yield the non-propagatable clones pDef-3p115 and pDef-3p91, respectively, both of which functioning as originally intended.

Targeted mutations in *vp4* were introduce by site-directed mutagenesis of p105 using o393+o394 and o395+o396 to yield p252 and p253, respectively. Golden Gate cloning of the latter plasmids in p57, yielded pDef-3p115-S and pDef-3p115-F, respectively.

#### Construction and modification of gene block plasmids

Plasmid pWT-noITR, pNS and pVP-3p115 have been published previously under the names pJA (18), pJΔVP and pJΔNS (19), respectively. Plasmid pVP served to produce pVP-3p49 and pVP-3p75 by site-directed mutagenesis, using o223+o227 and o223+o285, respectively. The same strategy was used to introduce 7m and 12m mutations in pNS (pVP-3p115) with o295+o296 and o297+o298, yielding pNS-7m and pNS-12m, respectively (pVP-7m and pVP-12m).

#### Construction of a replication reporter

AgeI and SacI were used to digest pDef-nf and pFAB217 (20). pDef-nf reaction products were dephosphorylated using antarctic phosphatase and ligated to pFAB217’s fragments using T4 DNA ligase to yield p73, in which the *ns* gene block is replaced by a small ~500 nts fragment of non-functional DNA. The PA3-eGFP cassette from pITR-A3-GFP (13) was amplified with o233+234. A large non-functional DNA fragment was amplified from pGC4783 (21) using o235+o236. These two amplicons were assembled by PCR using o233+o236, previously phosphorylated with T4 PNK. The resulting amplicon was cloned at the BbsI sites of p73. This yielded the replication reporter pITR-R, in which a reporter fragment of the same length as the original JcDV sequence is flanked by promoter-less ITRs.

### Cell line and transfection

The insect cell line Ld652Y (22) was maintained at 28°C in TC100 medium (Lonza) supplemented with 10% heat-inactivated fetal calf serum (Thermo scientific), and antibiotic/antimytotic cocktail (10 units/mL penicillin, 10 µg/mL streptomycin and 25 fg/mL Amphotericin B, Gibco). Ld652Y cells were originally derived from *Lymantria dispar* ovaries and have been found to be most sensitive to JcDV infection (18). Flask-grown cells at ~70% confluence were harvested, seeded on 24-well plates (10^5^ cells/well) and cultured overnight. Cells were then transfected with plasmids DNA (250 ng per plasmid per well) using FuGENE HD (Promega, 1:8 ratio), and cultured at 28°C. The culture medium was changed to a fresh one without transfection material 16h after transfection.

### Quantification of viral genome replication in transfected cells by qPCR

Transfected cells were resuspended in PBS at either 16h or 72h post-transfection and frozen in microtubes at −80C. Samples were lyzed through three freeze-thaw cycles (−80C/+20C) and clarified at 10,000 rpm for 10 minutes. For each sample, supernatants were diluted to a 100-fold in TE and 2.5 µL were directly used as templates in 10 µL qPCR reactions, using SYBR Green qPCR mix (Bioline) on a LightCycler 480 instrument (Roche). Each sample was assayed in technical triplicate, and their average used for further calculations. Primers o182+o183 (*neo*) were used to specifically quantify replication of the rearranged non-propagatable genomes, with an hybridization temperature of 54°C (Figure 2 and 6; Supplementary Figure 3). Primers o307+308 (*eGFP*) were used to quantify the replication reporter pITR-R, with a hybridization temperature of 60°C (Figure 3). The following amplification parameters were used: 2 min at 50°C and 2 min at 95°C, followed by 40 cycles each consisting of 15 s at 95°C and 30 s at the adequate hybridization temperature. Data were analyzed with LightCycler 480 software (version 1.5). Standard curves were generated after 40 cycles using ten 2-fold serial dilutions of p58. In each case, a linear range was obtained over the eleven standard points, enabling calculation of viral DNA copy number. Three biological replicates were performed for each assay.

**Figure 2.**
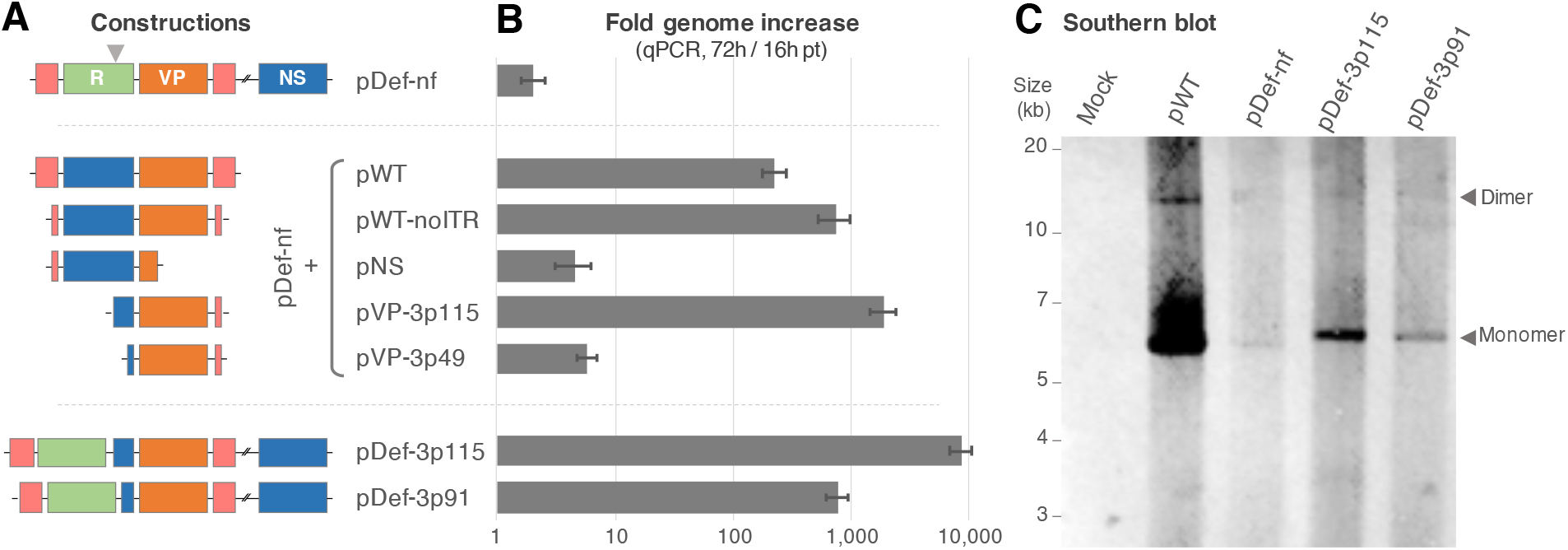
Impaired replication of non-functional clone is rescued by *vp* complementation. **A.** Schematic diagram of genetic constructions. ITRs are shown in red, *vp* in orange, ns in blue and reporter cassette in green. Sizes are not to scale. **B.** Global qPCR measurement of viral replication pinpoints region necessary and sufficient for rescue. Shown are fold changes in genome quantity between 16h and 72h post-transfection of Ld652Y cells with constructs depicted in A (qPCR target site marked by a grey arrowhead). The original design of the non-propagatable genome (top, pDef-nf) shows an almost complete absence of replication. Co-transfections with both the WT genome (pWT) and an ITR-less derivative (pWT-noITR) complement this phenotype, while a fragment restricted to the *ns* gene block does not (pNS). A large *vp* block comprising 115 nts downstream of the *vp* coding sequence (pVP-3p115) effects high rescue level, while one restricted to 49 nts of downstream context (pVP-3p49) does not rescue at all. Extension of *vp*’s native downstream context from 16 (pDef-nf) to 91 nts (pDef-3p91) or 115 nts (pDef-3p115) yields increasingly more functional non-propagatable clones that function as originally intended (Supplementary Figure 2). **C.** Apparent stalling of rolling-hairpin replication. Shown is a Southern blot probing *vp* sequences on DpnI-treated DNA extracted 96h post-transftection of Ld652Y with the indicated plasmids. Monomer-length and dimer-length species that are typical rolling-hairpin intermediate products are indicated by arrowheads. pDef-nf produces faint but detectable levels of intermediates. The functional derivatives pDef-3p115 and pDef-3p91 support elevated replication levels, but do not restore wild type performance. The discrepancy in relative performances of pWT between panels B and C is only apparent: panel B reports rescue of pDef-nf replication by pWT, while panel C reports replication of the pWT clone itself.

**Figure 3.**
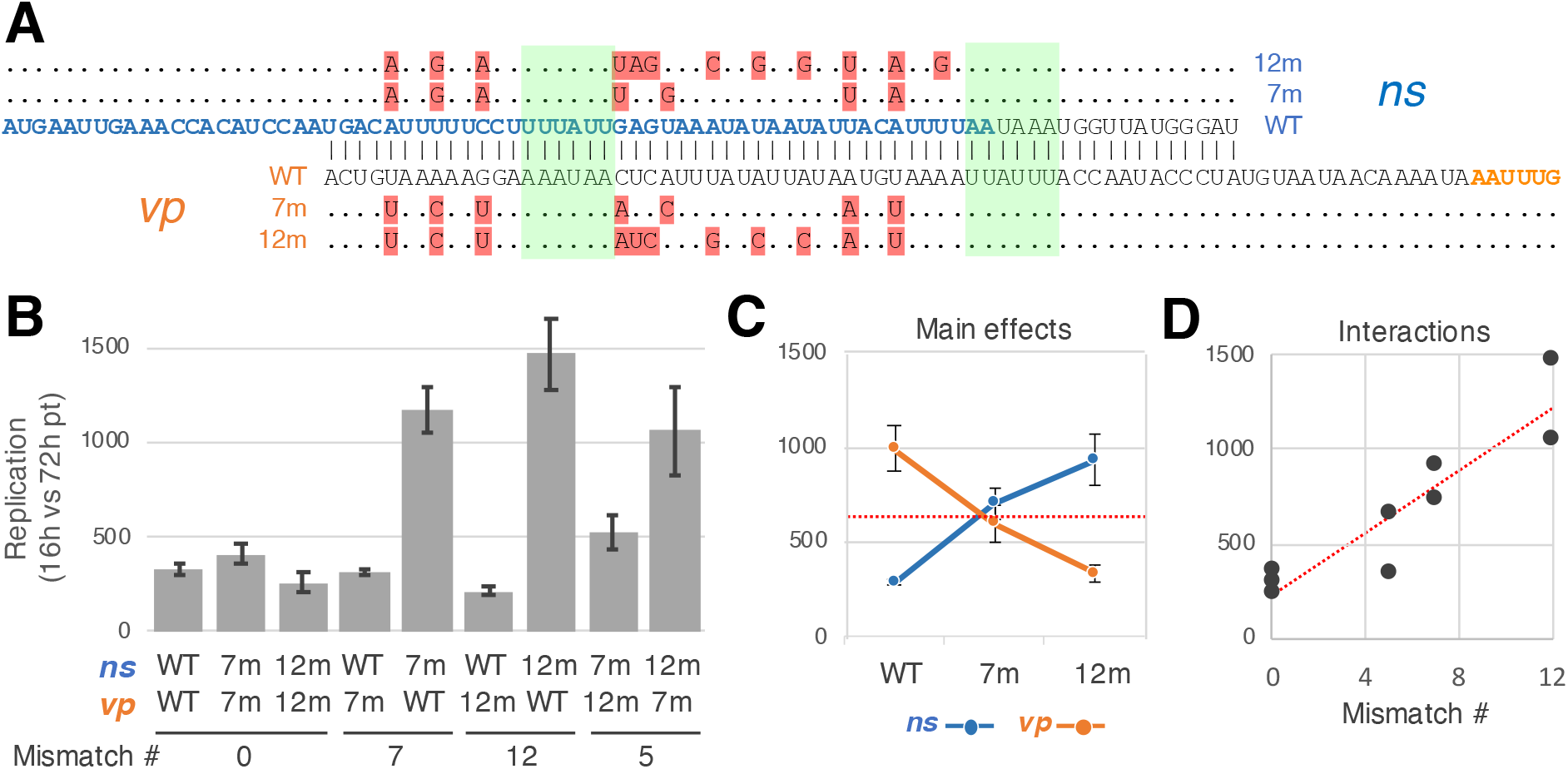
Antisense transcript interactions do not explain *vp*’s impact on replication. **A.** Alignments of WT and mutated transcripts in the overlapping region between *ns* and *vp*. WT *ns* and *vp* (reversed) transcript sequences are shown in the middle with ambisense complementarity marked by vertical bars. *ns* and *vp* coding sequences are colored in blue and orange, respectively. Complementary mutations (7 or 12) were independently introduced in distinct *ns* and *vp* constructs. Mutations (highlighted in red) correspond to synonymous *ns1* substitutions and avoid putative polyadenylation signals (green). Dots convey sequence identity. **B.** Mutations in *ns* and *vp* transcript show moderate but complex effects on replication. Shown are fold change in genome abundance as measured by qPCR between 16 and 72h post-transfection of Ld652Y cells by the indicated plasmid pairs and the replication reporter pITR-R. Error bars show standard errors across 3 biological replicates. While complementary mutations have little net impact on replication, mismatches show largely different effects on *ns* or *vp* transcripts. **C.** Contrasted effects of complementary mutations on *ns* and *vp*. Shown is the marginal mean fold change in genome abundance over the 3 assays involving each construct. Error bars show standard errors over 9 biological replicates. In line with antisense regulation of *ns*, mutations affecting the *ns* transcript tend to increase replication. Symmetrical mutations in the *vp* transcript show opposite effects, hinting at a distinct mechanism. **D.** Transcript mismatches modulate mutational effects. Shown are absolute values of interactions between constructs against the number of mismatches between *ns* and *vp* transcripts. Interactions represent the difference between measured fold changes and linear predictions based on main mutation effects plotted in panel C. Modulation by transcript mismatches confirms a role for antisense regulation.

Since there is no detectable virions released 72 hours post-transfection and genomes from intracellular virions are released from capsids by the thermal shocks, this assay quantifies transfected plasmids as well as all intermediates and final replication products present in the cells. The 16h time point quantifies the transfected plasmids before any sizable replication can occur (19). The ratio of measured genome quantities at 72h over 16h thus specifically quantifies replicated materials in intermediates or final forms.

### Quantification of viral transcripts abundance in transfected cells by RT-qPCR

Total RNA was extracted from cells 12h post-transfection using the RNeasy Mini kit (Qiagen). RNA samples were treated with Turbo DNAse (Ambion) and cDNAs were synthesized from polyA tails using SuperScript III reverse transcriptase (Invitrogen). Transcript abundances were measured with o40+o41 and o365+o366, which respectively binds to *vp4* and *ns1*, as described previously (19, 23). Cellular β-actin was used to control for RNA degradation and other potential sample to sample variations, as previously described (19). Each sample was assayed in technical triplicate and their average used for further calculations. Three biological replicates were performed.

### 3’RACE assays

Total RNA was extracted from cells 72h post-transfection using the RNeasy Mini kit (Qiagen). We used 5’/3’ RACE Kit, 2nd Generation (Roche) according to the manufacturer’s instructions. To specifically target *vp* transcripts we used primers o40 and o37 for the first and second round of PCR, respectively. Amplicons were Sanger-sequenced using o37. Chromatographs were plotted using ApE (v2.0.61).

### Quantification of protein abundance by immunofluorescence microscopy

Cells were transfected as described above and fixed 96h post-transfection with 4% paraformaldehyde for 20 min. Cells were washed twice in PBS and permeabilized with PBS+0.1% Triton (PBT). Blocking was performed with PBT+1%BSA for 1 hour. Cells were incubated with appropriate primary antibodies with PBT+0.1% BSA for 1 h, as described previously (19). Samples were washed twice in PBS and incubated overnight at 4°C in PBS. Samples were then incubated for 1h with anti-rabbit Alexa Fluor 568 secondary antibodies (Molecular Probes-Invitrogen) diluted at 1:500. Nuclei were labeled for 20 min with Hoechst 33342 (Sigma). Brightfield and fluorescence images were automatically acquired using a Cellomics platform equipped with 349 and 540 nm LED, Hoechst and Cy3 filter cube sets, a 10X Plan Apochromat 0.45NA objective and ORCA ER Hamamatsu digital camera. Images were batched processed with the Cellomics Studio software. Cell nuclei were automatically segmented using Hoechst signal. Abnormal signals were excluded to avoid artefacts. Mean Alexa Fluor 568 fluorescence signal per pixel in the segmented areas was computed for each cell in each sample.

### Western blots

Cells were seeded in 6-well plates (5.10^5^ cells/well), transfected with 1 μg of plasmid DNA and harvested 96h after transfection. Protein concentration was assessed by Bradford analysis, samples were diluted 1:20 and equal amounts of lysates were separated onto 4–15 % gel to run SDS-PAGE. Migrated materials were transferred to PVDF membranes (Immobilon-P, Millipore) for Western blot analysis. Anti-VP, -NS1, -NS2 (19) and anti-α-Tubulin (T9026, Sigma Aldrich) antibodies were incubated overnight at 4°C, followed by a 2 hours incubation with corresponding Horseradish peroxidase-coupled secondary antibody. Signals were detected by enhanced chemiluminescence (Luminata Crescendo, Millipore).

### Southern blots

Cells were seeded in 6-well plates (5.10^5^ cells/well), transfected with 1 μg of plasmid DNA and harvested 96h after transfection. Viral genomes were extracted using a Quick DNA miniprep plus kit (D4068, Zymo Research) and eluted in TE (100 μL). This eluate was digested with DpnI for 1h at 37°C to selectively cleave the transfected plasmids. Extracted genomes were resolved by slow overnight migration on a 0.8% agarose gel (25V), DNA was stained with ethidium bromide, blotted onto a nylon membrane (Roche Diagnostics, Indianapolis, Ind.) and fixed by UV irradiation (312 nm). Hybridization probes were an equimolar mix of PCR products amplified from 3 distinct *vp* regions using primers o405+o406 (784 bp), o407+o269 (896 nts) and o408+o409 (1072 bp). Labeling was achieved by using a 1:1 molar ratio of dCTP:Biotin-11-dCTP (NU-809-BIOX, Jena Bioscience) in the dNTPs used for the PCR (GoTaq DNA polymerase, Promega). The fixed membrane was incubated overnight at 68°C in presence of the hybridization probes and revealed using a biotin chromogenic detection kit (K0661, Thermofischer Scientific).

### Data analysis

GT-content analysis was performed using custom python script and plotted using R. The factorial analysis presented in Figure 3 was performed using the *anova* function in R. Analysis of Cellomics data used custom R scripts. Scripts and data are available upon request to GC

## Results

### Translocation of ns gene cluster abolishes replication of a rearranged JcDV genome

In parvoviruses, replication occurs through a rolling-hairpin mechanism that involves *ns* gene functions and the terminal hairpins that flank the genome (ITRs; Figure 1A; (24). These telomeric regions are also important for genome packaging. In an effort to construct a safe, non-propagatable variant of a JcDV infectious clone, we replaced the *ns* gene block by a reporter cassette and cloned it under a strong heterologous promoter on the same plasmid, but outside of the viral genome, as delineated by the telomeres (pDef-nf, Figure 1B). We expected that, upon transfection, *ns* genes would be expressed at sufficient levels to ensure replication of the modified genome from its telomeres and its subsequent packaging in capsids originating from *vp* genes. To avoid any size-related packaging problem (25), the reporter cassette was designed to be the same size as the translocated *ns* block. Cells transduced by the resulting virions would produce VP proteins, a drug resistance protein and a fluorescence protein, but would not be able to support further replication or packaging in the absence of NS functions (Figure 1CD).

The terminal hairpins that flank the genome of parvoviruses are particularly refractory to PCR amplification. This has typically restricted modifications of these genomes to those permitted by available restriction sites. Here, we developed a Golden Gate cloning strategy to enable more precise manipulations, and could effortless and seamlessly assemble the designed plasmid above from individual building blocks (see Materials and Methods and Supplementary Figure 1). This plasmid could be transfected with an efficacy of about 60%, as estimated by counting fluorescent cells 3 days post-transfection (pt). JcDV’s cycle is quite long compared to other parvoviruses, and virions are released from bursting cells 5-7 days after infection or transfection. Upon transduction of naïve cells with a supernatant of the above culture collected at 7 days pt, we could not observe any fluorescent cell, indicating that the pDef-nf construct did not behave as expected (Supplementary Figure 2). All JcDV proteins for which we possess antibodies (VP, NS1 and NS2) could be qualitatively detected by immunofluorescence upon transfection of the plasmid (data not shown), suggesting that the defect did not originate from complete absence of expression. Quantification of viral genome abundance after transfection, however, demonstrated an almost complete absence of replication (Figure 2, top).

### Complementation assays link replication defect to vp gene

As NS proteins are involved in replication, we reasoned that the translocation of the *ns* block could negatively impact its expression pattern. We expected that viral genes moved outside of the genome would experience a relative decrease in gene dosage, as the viral genome is replicated. In our design, we had sought to compensate this by using the immediate early 2 promoter from the multicapsid Nuclear Polyedrosis Virus of *Orgyia pseudotsugata* (POpIE2)—one of the strongest heterologous insect promoter (26)—to drive *ns* expression. As insufficient expression of NS functions would effect inefficient replication rather than its complete shutdown, we conjectured that the observed defect could rather stem from an ineffective expression pattern linked to the absence of appropriate regulatory signals in the heterologous promoter sequence. To explore this hypothesis, we tested three different constructs in which POpIE2 was replaced by *i)* the 185 nucleotides (nts) immediately upstream of *ns3*’s start codon (core P93 promoter); *ii)* a larger fragment spanning 333 nts upstream (extended P93 promoter); and *iii)* a larger fragment of 595 nts (full upstream sequence missing the left end of the terminal hairpin). None of these modifications remediated the impaired replication phenotype.

We next used a genetic complementation strategy to determine whether the non-functional pDef-nf clone could be rescued in *trans* (Figure 2A). Co-transfection with the infectious JcDV clone was indeed able to restore replication of pDef-nf, as measured by qPCR targeted to the engineered reporter cassette (Figure 2B, pDef-nf+pWT). Likewise, a construct in which ITRs have been clipped down to the core P9 and P93 promoters was capable of functional rescue (Figure 2B, pDef-nf+pWT-noITR). Since such a clone cannot replicate, this result disproves the potential implication of a replication-related disbalance in *ns* gene dosage. A truncated clone comprising only the *ns* gene block driven by its core promoter could not rescue replication (Figure 2B, pDef-nf +pNS), further indicating that the replication defect is not directly related to NS functions. Instead, we found that complementation with the *vp* gene block only was sufficient to restore replication (Figure 2B, pDef-nf +pVP-3p115). This result was particularly surprising, as capsid genes were not known to be involved in JcDV’s replication process.

The rescuing *vp* block differs from the sequence in the non-functional pDef-nf clone by 99 nts of natural sequence context downstream of the *vp* gene. We thus trimmed that region to test complementation by a *vp* gene block comprising only 33 additional nts, and found that it was not sufficient for rescue (Figure 2B, pDef-nf +pVP-3p49). To confirm and extend these results, we constructed two amended versions of the non-functional clone in which the *vp* block is extended by 99 and 75 nts, respectively (Figure 2B, pDef-3p115 and pDef-3p91). Both are capable of independent replication and represent functional non-propagatable clones that behave as originally intended (Figure 1D and Supplementary Figure 2).

We next performed Southern blots to more finely resolve the replication behaviors of these non-propagatable constructs (Figure 2C). In agreement with qPCR signals, we found that pDef-nf is only capable of producing very weak levels of replication intermediates, suggesting that the rolling-hairpin mechanism is functional but stalls in its first cycle(s). pDef-3p91 and pDef-3p115 showed incrementally higher levels of replication products—as expected from qPCR and functional data—but do not rival with the performance of the WT clone.

### Transcript antisense interactions may repress replication but do not explain the role of vp

Although the results above readily fixed the faulty original design, they brought forward an unexpected role for the sequence region downstream of the *vp* coding sequence. Consequent to its ambisense genome architecture, JcDV’s *ns* and *vp* transcripts antisense overlap by 60 nts on their respective 3’ sides, with *vp*’s 3’UTR overlapping the *ns1* coding sequence by 44 nts (Figure 1A). We noted above that parvoviruses’ terminal hairpins are not well processed by PCR polymerases. They are equally problematic for *E. coli*’s polymerase, which creates recombinogenic conditions that render plasmid clones unstable during propagation in these bacteria. This issue is particularly marked with JcDV’s long and stable hairpins (130 nts). In our original design, we had therefore opted to truncate *vp*’s 3’UTR by 59 nts in order to avoid introduction of recombination-prone sequence duplications (Figure 1B).

We reasoned that the absence of transcript overlap in pDef-nf could prevent replication if an antisense regulation mechanism was somehow involved in upregulating *ns* expression (5, 23). To investigate whether base-pairing of the two transcripts is necessary for replication of the viral genome, we adopted a complementation strategy that enable independent manipulation of the antisense region on each transcript. We produced two sets of variants with 7 and 12 substitutions that are both synonymous in the context of NS1 coding sequence and avoid alteration of putative poly-adenylation signals (Figure 3A). We then tested the effects of combining these sequences on the replication of a reporter construct.

Our results demonstrate sizable impacts of the substitutions, with up to 7-fold difference in replication efficiency (Figure 3B). However, the mechanism portrayed by our data is more complex than expected, with matching mutations displaying contrasted effects on the *ns* or *vp* blocks. Substitutions in *ns* are generally associated with increased replication (Figure 3C, light grey), unless they are complemented by matching *vp* mutations (Figure 2B), which suggests that antisense regulation normally limits the expression of *ns* genes. Nonetheless, antisense mismatches driven by corresponding substitutions in *vp* hardly impact replication (Figure 2B), supporting the notion that *vp* mutations exert some general, negative impacts on replication (Figure 3C, dark grey) that could compensate disruption of antisense regulation. Further supporting the existence of antisense regulation, we found that the number of mismatches between pairs of variants is associated with their statistical interactions (*i.e.* how much their joint impact is different from expectations based on their individual effect on replication, Figure 3D). Though more thorough investigations would be needed to unambiguously conclude on the existence of antisense regulation between *ns* and *vp* transcripts, it appears that this mechanism might play a role in decreasing viral replication’s level. Accordingly, the missing antisense region in our original pDef-ns construct would be expected to promote replication rather than impair it. We conclude that perturbation of potential antisense regulation is unlikely to be the cause of the observed replication stall.

### vp’s full 3’UTR is required for replication and determined by downstream sequence context

So far, we have shown that the 43 nts region extending from position +50 (pVP-3p49) to +91 (pDef-3p91) after *vp*’s stop codon is strictly required for replication (Figure 2). The most apparent functional features in this region are the polyadenylation (polyA) signal and site of *vp*’s transcript (Figure 1), respectively located at position +57:+62 and +75 (23). As mutating the *vp* transcript between these positions appears to negatively impact replication (Figure 3C), we sought to determine whether production of a complete *vp* transcript would be sufficient for rescue.

We constructed a *vp* block truncated just after the polyA site (pVP-3p75) and found that it is not capable of supporting viral replication, yielding qPCR readouts similar to that of the shorter pVP-3p49 in the context of our complementation assay (Supplementary Figure 3). This prompted us to perform 3’RACE to verify the exact position of *vp*’s polyA site. The polyA site was conform to previous report (23) in both the WT clone and the rescue-positive *vp* block pVP-3p115 (Figure 4A). In contrast, the transcript produced by the clipped-to-polyA-site pVP-3p75 block is shortened by 17 nts.

**Figure 4.**
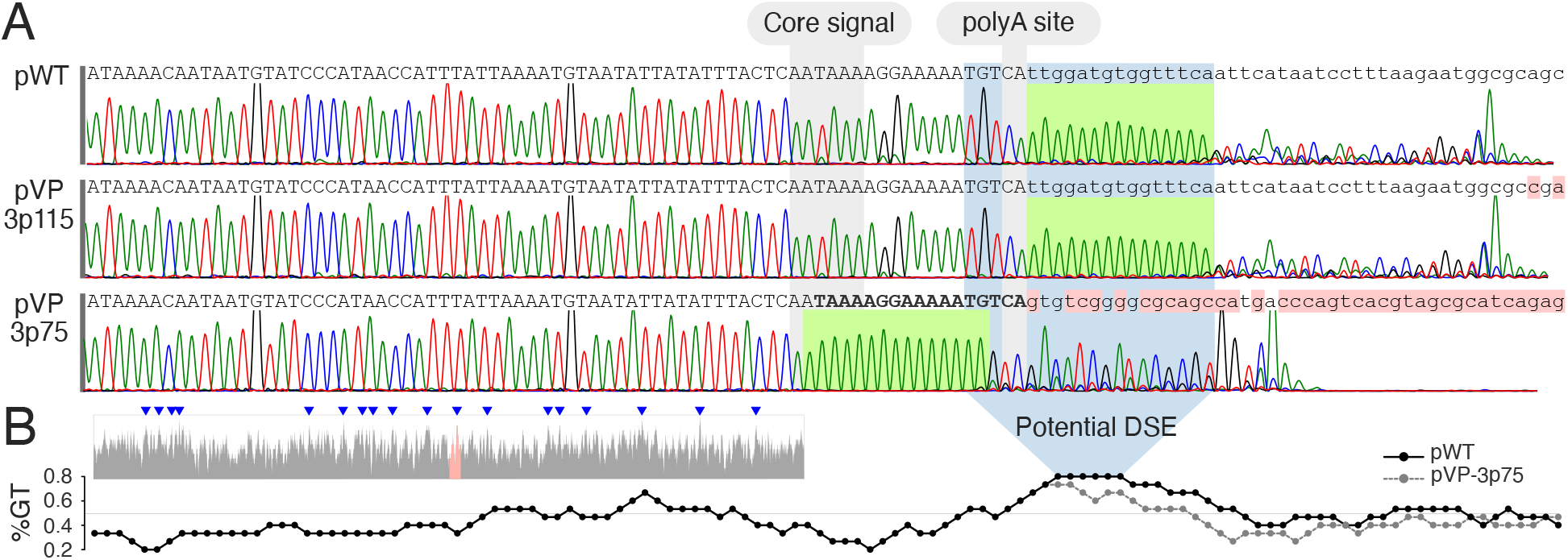
Full *vp* 3’UTR is required for replication and specified by its immediate downstream context. **A.** Impaired replication is associated with truncated 3’UTR. Shown are representative chromatograms of the sequence downstream of *vp*’s stop codon as obtained by 3’RACE. The sequence corresponding to each construct is shown on top. Mismatches (red background) correspond to the sequence context of the plasmid backbone. A construct comprising 115 nts of native context (pVP-3p115) produces the same transcript as the WT clone. A construct truncated just after the polyA site (pVP-3p75) produces shorter transcripts that lack the last 17 nts (bold) of the native 3’UTR and do not support replication (Supplementary Figure 3). **B.** Identification of a potential Downstream Sequence Element (DSE). The inset shows the %GT calculated for sliding windows of 15 nts over JcDV’s genome. Blue arrowheads mark the 18 peaks with %GT≥80. The red region corresponds to the sequence shown in panel A and is enlarged in the main plot, where points show %GT of sliding windows centered at these positions. The %GT peak located immediately downstream of the polyA site represents a potential DSE (solid black line) that is largely attenuated in pVP-3p75 (dashed grey line).

Molecular signals determining polyadenylation are known to be largely context dependent. In particular, T- or GT-rich downstream sequence elements (DSE) have been shown to play an important role in some instances (27). We thus carried a bioinformatic analysis of JcDV’s genome and identified a local %GT peak just downstream of *vp*’s polyA site that is diminished in the sequence context of pVP-3p75 (Figure 4B).

Altogether, these data show that production of a native *vp* transcript with full-length 3’UTR is essential for effective replication and might depend on a GT-rich downstream sequence to suppress a cryptic, upstream polyA signal.

### VP’s 3’UTR control VP but not NS expression

To further investigate the mechanistic link between *vp* transcript and replication, we sought to determine potential feedbacks of altered *vp* expression on *ns* expression. We first quantified the abundance of *ns* and *vp* transcripts using reverse transcription quantitative PCR (RT-qPCR). To avoid confounding gene dosage effects linked with differential replication, samples were collected 12h after transfection *i.e.* before any sizable replication can be detected (19). Cells transfected by the non-functional version of the non-propagatable clone (pDef-nf) show a 100-fold decrease in *vp* transcripts as compared to cells transfected with an amended functional version (pDef-3p115). In sharp contrast, *ns* transcript abundances are identical in both background (Figure 5A). We conclude that truncation of *vp* transcript’s 3’UTR results in its markedly decreased abundance, but does not appears to impact the *ns* transcript.

**Figure 5.**
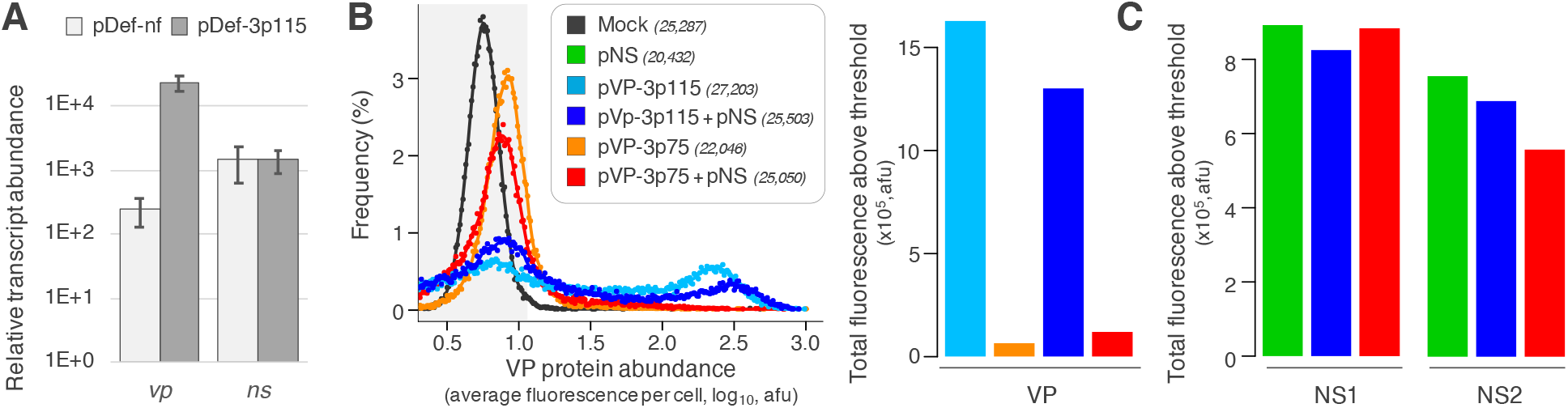
Alteration of *vp*’s 3’UTR impacts VP but not NS expression. **A.** Shorter 3’UTR decreases *vp* transcript abundance. Shown are relative transcript abundances, as quantified by RT-qPCR 12h after transfection of Ld652Y cells by the indicated plasmids. A functional version of the non-propagatable clone (pDef-3p115, dark grey) produces native *vp* transcript that cumulates to a 100-fold higher level than truncated transcripts from the non-functional clone (pDef-nf, light grey). Abundances of *ns* transcripts are identical for both constructs. **B.** Shorter 3’UTR decreases VP protein abundance. Distributions of cellular fluorescence intensities (left), as quantified by immunofluorescence microscopy for VP proteins 3 days after transfection of Ld652Y cells by different combination of *vp* and *ns* blocks, as color coded in the legend. The number of analyzed cells is indicated in parentheses. Data points represent cell frequencies for each percentile of the fluorescence range. For visual clarity, points are underlaid with loess regression smoothers. The 99^th^ percentile of control cells (grey background) is used as a threshold to quantify positive fluorescence signals. Barplot (right) shows the sum of fluorescence intensities for cells above threshold. Consistent with lower abundancies (panel A), truncated *vp* transcripts lead to strongly reduced VP protein level, both in the absence and presence of *ns* genes. **C**. Reduced VP expression has little impact on NS expression. Barplots of total fluorescence intensities calculated as in panel B for cells marked for NS1 and NS2, respectively (Supplementary Figure 4AB). Color code as in panel B. VP is not involved in upregulating NS expression.

We next used immunofluorescence microscopy to quantify protein abundances and identify potential post-transcriptional regulations. To avoid confounding replication feedback on gene dosage while allowing sufficient time for protein synthesis, we co-transfected cells with combinations of non-replicative *ns* and *vp* blocks and quantified consequent VP, NS1 and NS2 signals 3 days post-transfection. Consistent with RT-qPCR data, we found that a construct yielding truncated *vp* transcripts leads to markedly lower VP protein abundances (Figure 5B, pVp-3p75). Co-transfection with the *ns* block results in a modest 1.8-fold increase in overall VP signal from that pVp-3p75 and in a slight 1.3-fold decrease for the functional pVp-3p115 (Figure 5B, right).

Nonetheless, high VP-producing cells in the latter are clearly shifted toward higher production, with a 1.4-fold increase in the higher distribution mode (Figure 5B, left). This discrepancy might reflect differential co-transfection efficiencies. In any case, the presence of NS proteins does not appear to have a dramatic impact on VP expression. Similarly, our data does not suggest a strong impact of either native nor non-functional *vp* blocks on the expression levels of NS1 or NS2 (Figure 5C). We confirmed these results using Western blots (Supplementary Figure 4C), and could further observe that both versions of the functional non-propagatable genome (pDef-3p91 and pDef-3p115) could express VP, with a small increase in NS1 abundance with respect to the non-functional pDef-nf (Supplementary Figure 4D).

Altogether, our results show that neither the nature of *vp*’s transcripts nor their differential consequences on VP protein abundances have a strong effect on the expression of *ns* genes. Thus, the impact of VP’s 3’UTR on replication does not appear to involve any theretofore unknown feedback on NS regulation at either transcriptional and translational level. Instead, our data strongly suggest a more direct and essential role in replication for VP proteins, which is prevented by markedly lower protein expression from transcripts with truncated 3’UTR. To further verify this, we constructed two variants of the functional non-propagatable clone pDef-3p115. In the first, we substituted a nucleotide at position 1,626 of the 2,433 nts-long *vp* coding sequence to introduce an Amber stop codon (pDef-3p115-S). In the second, we deleted a nucleotide at position 2,061 to produce a frameshift (pDef-3p115-F). Both constructs produce truncated VP proteins (Supplementary Figure 4D). The effect of these truncations on replication is largely similar to that of a non-functional 3’UTR, as evidenced by qPCR measurements and Southern blots (Figure 6). Since these mutations entail minimal differences with WT sequence and are separated by more than 400 nts, it is highly unlikely that their effects are mediated by anything else than their impact on translation. We found no other ORF than *vp* spanning the extent of these two locations on either strand. Therefore, these results strongly suggest that the VP proteins themselves exert the observed impact on JcDV’s replication. This *trans* effect does not appear to prevent the onset but the continued processivity of the rolling-hairpin replication mechanism, as we observed low levels of replication intermediates for the truncated constructs (Figure 6C).

**Figure 6.**
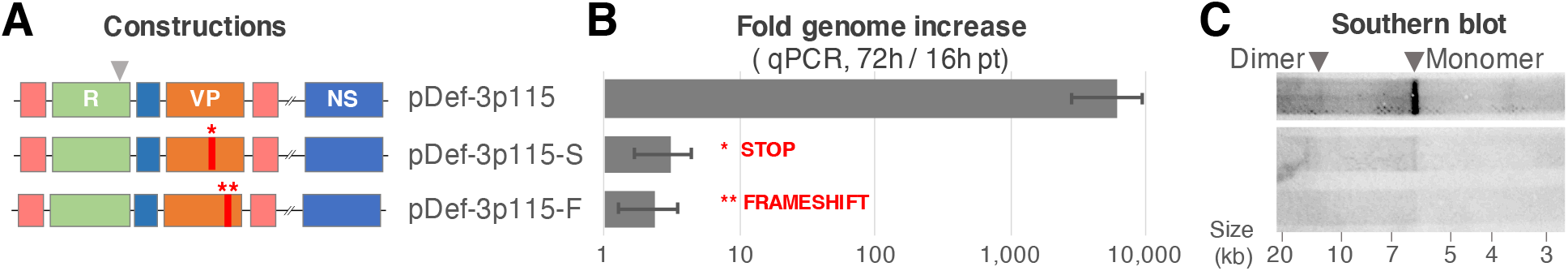
VP proteins are directly involved in the replication process. **A.** Schematic diagram of genetic constructions. ITRs are shown in light red, *vp* in orange, ns in blue and reporter cassette in green. Red bars and asterisks mark the location of mutations resulting in the production of truncated VP proteins (Supplementary Figure 4D). Mutations correspond to a stop (*: TA**T** 542 TA**G**; pDef-3p115-S) or a frameshift (**: AA**T** 687 AA-; pDef-3p115-F). Sizes are not to scale. **B.** Truncating VPs proteins abolishes viral replication as measured by qPCR. Shown are fold changes in genome quantity between 16h and 72h post-transfection of Ld652Y cells with constructs depicted in A (qPCR target site marked by a grey arrowhead). Data for pDef-3p115 are reported from Figure 2B for comparison. **C.** Apparent stalling of rolling hairpin replication by truncated VPs. Shown is a Southern blot probing *vp* sequences on DpnI-treated DNA extracted 96h post-transftection of Ld652Y with the plasmids depicted in A. Monomer-length and dimer-length species are indicated by arrowheads. Both pDef-3p115-S and pDef-3p115-F produce faint but detectable levels of intermediates. Lane for pDef-3p115 is reported from Figure 2B. It originates from the same image with identical manipulations and is directly comparable. Since the introduced mutations involve minimal DNA sequence modification and are separated by 435 nts, these data strongly support the link between VP proteins and replication.

## Discussion

We developed an efficient Golden Gate cloning strategy to precisely alter otherwise hard-to-manipulate parvovirus genomes (Supplementary Figure 1). We used this approach to genetically dissociate the capsid and non-structural gene clusters of the insect parvovirus JcDV (Figure 1). To implement this genomic rearrangement, we initially chose to truncate the 3’UTR of the capsid gene, which naturally overlaps the non-structural cluster to be relocated. This led us to uncover a profound impact of the capsid gene products on the replication of this parvovirus (Figures 2, 5 and 6).

We have shown that the region immediately downstream of *vp*’s polyA site is necessary to suppress an upstream polyA signal. This region likely comprises a GT-rich Downstream Sequence Element (DSE) that modulate polyadenylation (Figure 4B). Although more work is required to delineate the exact boundaries of this element, our data suggest that it functions in a cumulative fashion, with longer downstream sequences associated with stronger effects on VP expression levels and genome replication (e.g. pDef-3p115 vs pDef-3p91, Supplementary Figure 4D and Figure 2BC).

In the absence of the putative DSE, a cryptic polyA site yields an alternate transcript species that is 17 nts shorter than the WT (pVP-3p75, Figure 4A). Such shorter 3’UTRs determines markedly decreased *vp* transcript abundances and VP proteins production (Figure 5AB and Supplementary Figure 4CD), and are associated with an almost complete absence of viral genome replication (Figure 2BC and Supplementary Figure 3). The last 17 nts of the native 3’UTR may contain binding motifs for proteins that increase transcript stability, thus enabling higher protein production from more abundant transcripts. Alternatively, regulatory proteins binding to such motifs could promote translation through RNA looping, and increased transcript stability would arise as a consequence of more active translation (28). Known binding motifs could not be detected in this region (29). Instead of sequential motifs, some RNA binding proteins recognize structural signals (30). *In silico* prediction shows that the end of the native 3’UTR can fold into a convincing secondary structure that exhibit a central bulge typical of such signals (31) Supplementary Figure 5A). The lower stem of that structure cannot form in the truncated transcript, due to the deletion of the corresponding sequence (Supplementary Figure 5B). In *vp*’s 3’UTR variants for which we identified potential negative impacts on replication (7m and 12m, Figure 3), the cryptic polyA site is mutated and the predicted secondary structure is weakened to various extents (Supplementary Figure 5CD). Although regulatory proteins known to bind secondary structures in the 3’UTR usually inhibit translation, the structure at the end of *vp*’s transcript could be involved in translation activation or, possibly, increased stability. Regardless of the actual mechanism, the decrease in VP protein production due to premature transcript termination causes a dramatic and unexpected stalling of viral replication.

In the context of the natural JcDV genome sequence, we found no evidence for strong *trans* expression feedbacks between VP and NS proteins, as has been documented for other parvoviruses (32–36). In particular, the impact of decreased abundance in *vp* transcript and VP proteins on replication does not appear to involve a direct effect on NS1 or NS2 (Supplementary Figure 4ABC and Figure 5C). While the precise role of NS2 is unknown, NS1 possesses nickase and helicase activities that are essential for rolling-hairpin replication (24, 37). Although NS3 is also known to be essential for JcDV’s replication, we could not investigate its expression pattern in this study for an effective cognate antibody is lacking (38). Nonetheless, we surmise that NS3’s expression is highly unlikely to be impacted by VP. Indeed, both NS1 and NS2 are translated from transcripts spliced of *ns3* in such a way that all these genes present an almost identical 5’UTRs (Figure 1A; (23). Since VP proteins have little effect of the production of NS1-2, it is reasonable to infer that they do not impact splicing and thus have no effect on any splicing- nor 5’UTR-related regulations of *ns3*’s expression.

In the context of the engineered, non-propagatable clones—in which the *ns* block is located outside of the genome—we observed that enabling VP expression and consequent viral replication is associated with a moderate increase of NS1 abundance (pDef-nf vs pDef-3p115 and pDef-3p91, Supplementary Figure 4D). In contrast, expression of truncated VP proteins that do not support replication appears to be associated with lower abundances of both NS1 and NS2 (pDef-3p115-S and pDef-3p115-F, Supplementary Figure 4D). VPs could possibly interact with the heterologous promoter used to drive *ns* expression in these constructs, as was recently shown for AAV capsids (39). This, however, would be highly unlikely given that the steady-state level of *ns* transcripts is not impacted by changes in VPs’ abundances (Figure 5A). A context-specific impact of VPs on NS translation is also difficult to conceive. Rather than an effect on expression, these variations could reflect differential stability of NSs, perhaps mediated by interactions with VPs that depends on effective replication.

Our data suggest an essential role for VP proteins in the replication process of JcDV (Figure 6), through a mechanism that involves either protein interactions with NS proteins or autonomous replicative function for isolated VP proteins, their capsomeres or entire capsids. A few reports on other parvoviruses have pointed in these directions. For example, it was shown that combined mutations in VP2 of the mink enteritis parvovirus can diminish genome replication by up to 10 fold (40). A similar decrease in replication efficiency was also associated with VP2 mutations in porcine parvovirus (PPV; (41). As potential impacts on *ns* regulation were not tested in these studies, it is however difficult to conclude on the exact role played by these VP proteins.

More interestingly, replication of Adeno-Associated Virus Type 2 (AAV-2) and derivative recombinant genomes used in gene therapy was shown to stall to very low levels in the absence of efficient packaging (42). Further, both encapsidation and elongation levels were found to be higher when the capsid gene (*cap*) is provided in *cis* rather than in *trans* of the packaged genome. Although genetic systems supporting high replication in the absence of capsid proteins have been described, these observations suggest that a tight spatio-temporal coupling between replication, capsid protein expression and encapsidation improves the natural productivity of this virus. It was later elegantly confirmed that the formation of AAV-2 virions preferentially involves capsid proteins that are physically produced from the very genome that is packaged (11). Similar couplings have been described for positive-strand RNA viruses (43). Although the exact mechanistic basis of this process remains unknown, our observations that deficient VP expression or truncated proteins stall replication and can be partially rescued by *trans* complementation are compatible with such a phenomenon. It is thus tempting to speculate that the dependency of replication over packaging could be a shared feature of parvoviruses. The magnitude of the phenotypes reported here suggests that JcDV provides an excellent model to further probe this mechanism.

All icosahedral viruses define a limited inner capsid volume that impose strong constraints on the upper size of the packaged genome (44, 45). From an evolutionary perspective, these constrains can be accommodated either physically—by increasing the capsid triangulation number to fit a larger genome or parting the latter into multiple capsids (multipartism); or informationally— through genetic information compression (*e.g.* overlapping genes). While multipartism could participate in increasing viral modularity (46), informational compression might create functional dependencies between seemingly independent blocks that could greatly diminish it. Our observation that some function coded in the *vp* block is absolutely required for viral genome replication initially led us to question the effective modularity of JcDV’s architecture, and more broadly of other parvoviruses and viruses. Our investigations did not permit to fully resolve the precise mechanism underlying this dependency. If, as suggested above, this mechanism involves efficient packaging of the replicating genome rather than very specific interactions with NS proteins, that dependency may not in fact compromise modularity, nor its supposed evolutionary benefits in terms of evolvability. On the contrary, such a mechanism would prevent genome variants that are unable to produce propagatable capsid from depleting cellular resources via replication at the expense of other functional variants. This would also limit their hitchhiking functional capsids and enforce the genetic linkage between capsid variants and their corresponding genomes, thus facilitating the evolution of functional capsid variants.

## Supplementary data

Supplementary Figures 1–5 and Supplementary Tables 1–2 are available with the online version of this article.

## Acknowledgements

We are grateful to C. Clouet, M. Schatz and A.S. Gosselin-Grenet for useful discussions and technical assistance.

## Supplementary data

**Supplementary Figure 1.**
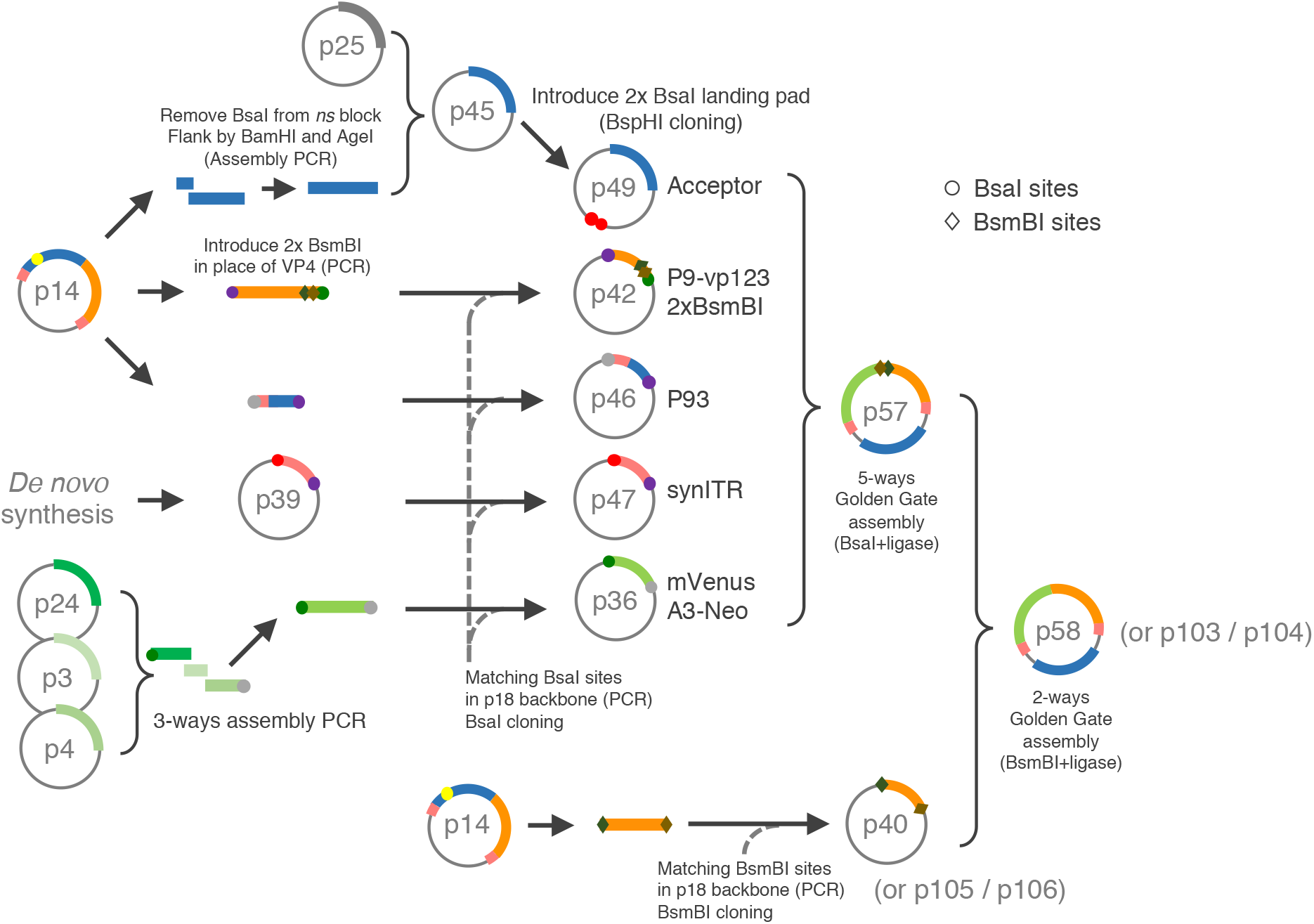
Efficient Golden Gate cloning strategy for precise modification of parvovirus genomes. The *ns* block—modified to erase a naturally occurring BsaI site—is cloned along with a double BsaI landing to form an acceptor plasmid (p42). Individual building blocks are flanked by BsaI sites by PCR, cloned in donor plasmids and sequence verified (p46, p47 and p36). A five-ways Golden Gate assembly of donor and accepting plasmid led to efficient assembly of the designed genome with a single reaction (p57). The order of assembled fragments is coded in the overhangs created upon BsaI digest (colored circles). Aside from the plasmid containing the PCR-refractory terminal hairpin (p39), all donor plasmids are easily amenable to modification by site directed mutagenesis. Because the original intent was to easily introduce mutation in *vp4*, this fragment was designed to be cloned in a final BsmBI Golden Gate cloning step into a *vp4*-less intermediate acceptor construct, yielding the final construct (p58, referred to as pDef-nf in the main text). The original *vp4* fragment (p40) was latter extended by site directed mutagenesis to increase the length of the 3’tail (p105 and p106) and yield functional non-propagatable clones (p103 and p104, referred to as pDef-3p115 et pDef3p91 in the main text, respectively).

**Supplementary Figure 2.**
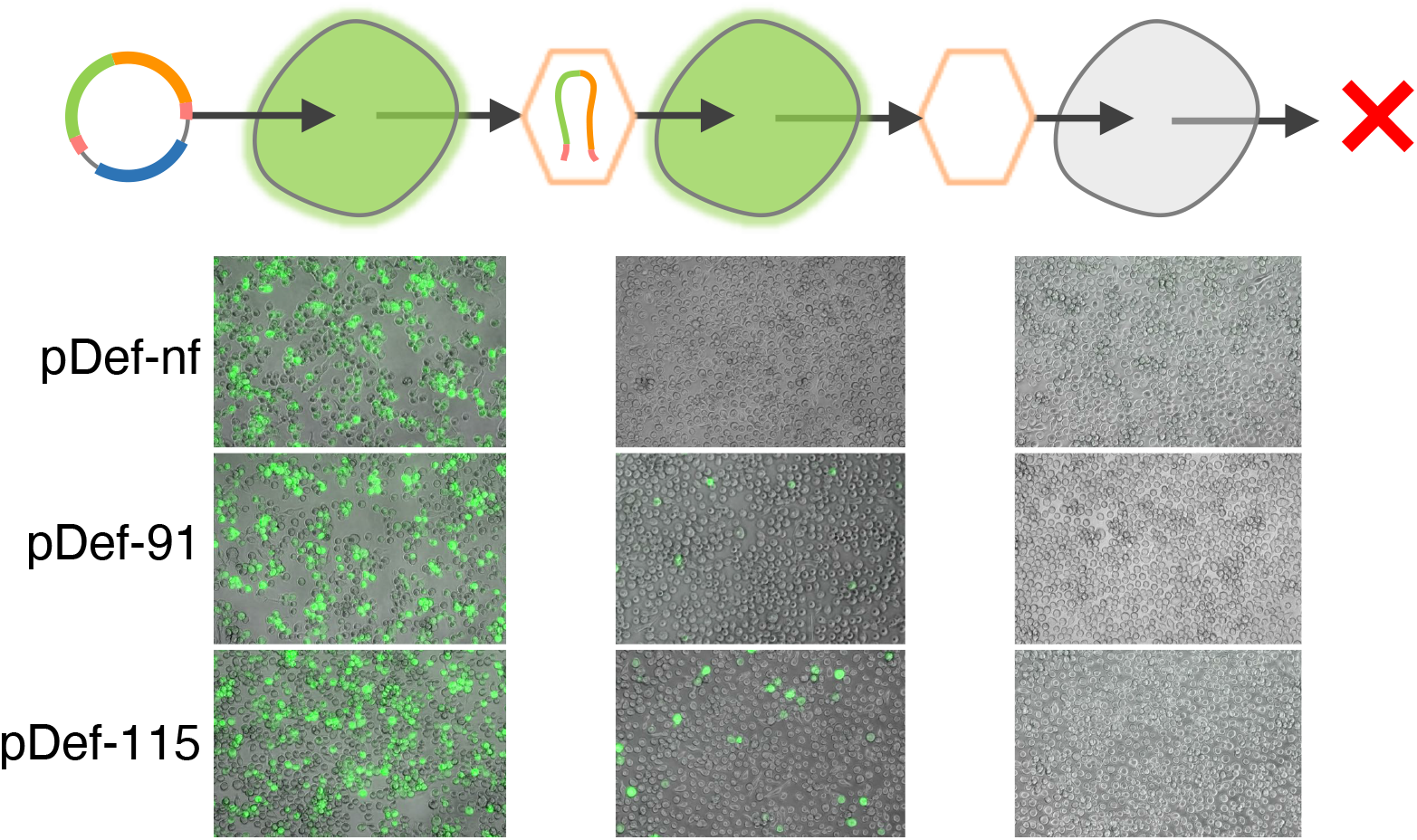
Amended versions of the non-propagatable clone function as intended. Images where taken using a regular DSLR camera fitted to an epifluorescence microscope. The first column shows Ld652Y cells 3 days post-transfection with different plasmids, as shown. The second column show naïve cells 3 days post-transduction with 100 μL of clarified supernatants obtained from lysing transfected cells by repeated heat shock at 7 days pt. The third column show naïve cells 3 days post-transduction with 100 μL of clarified supernatants obtained from lysing first-round infected cells by repeated heat shock at 7 days pt. While no fluorescence could be observed for the first round of infection with the non-functional pDef-nf construct, transduction was effective for the amended pDef-3p91 and pDef-3p115, to an extent commensurate with observed replication levels (see Figure 2BC). The second round of infection shows no signs of transduction, as expected from non-propagatable clones.

**Supplementary Figure 3.**
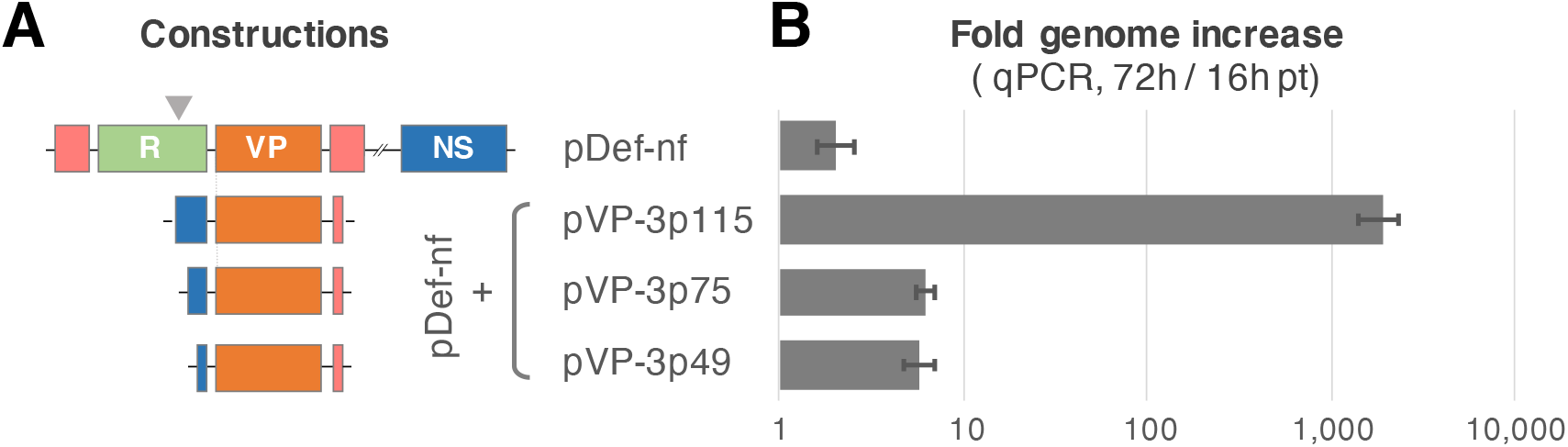
VP block truncated just downstream of the polyA site does not rescue replication. **A.** Schematic diagram of genetic constructions. ITRs are shown in red, *vp* in orange, ns in blue and reporter cassette in green. Sizes are not to scale. **B.** Global qPCR measurement of viral replication. Shown are fold changes in genome quantity between 16h and 72h post-transfection of Ld652Y cells with constructs depicted in A (qPCR target site marked by a grey arrowhead). Data for pDef-nf, pDef-nf+pVP-3p115 and pDef-nf+pVP-3p49 are reported from figure 2B for comparison. Construct pVP-3p75 corresponds to pVP-3p115 truncated just after the CA of the polyA site (see Figure 4) and yields very low replication level when complementing pDef-nf.

**Supplementary Figure 4.**
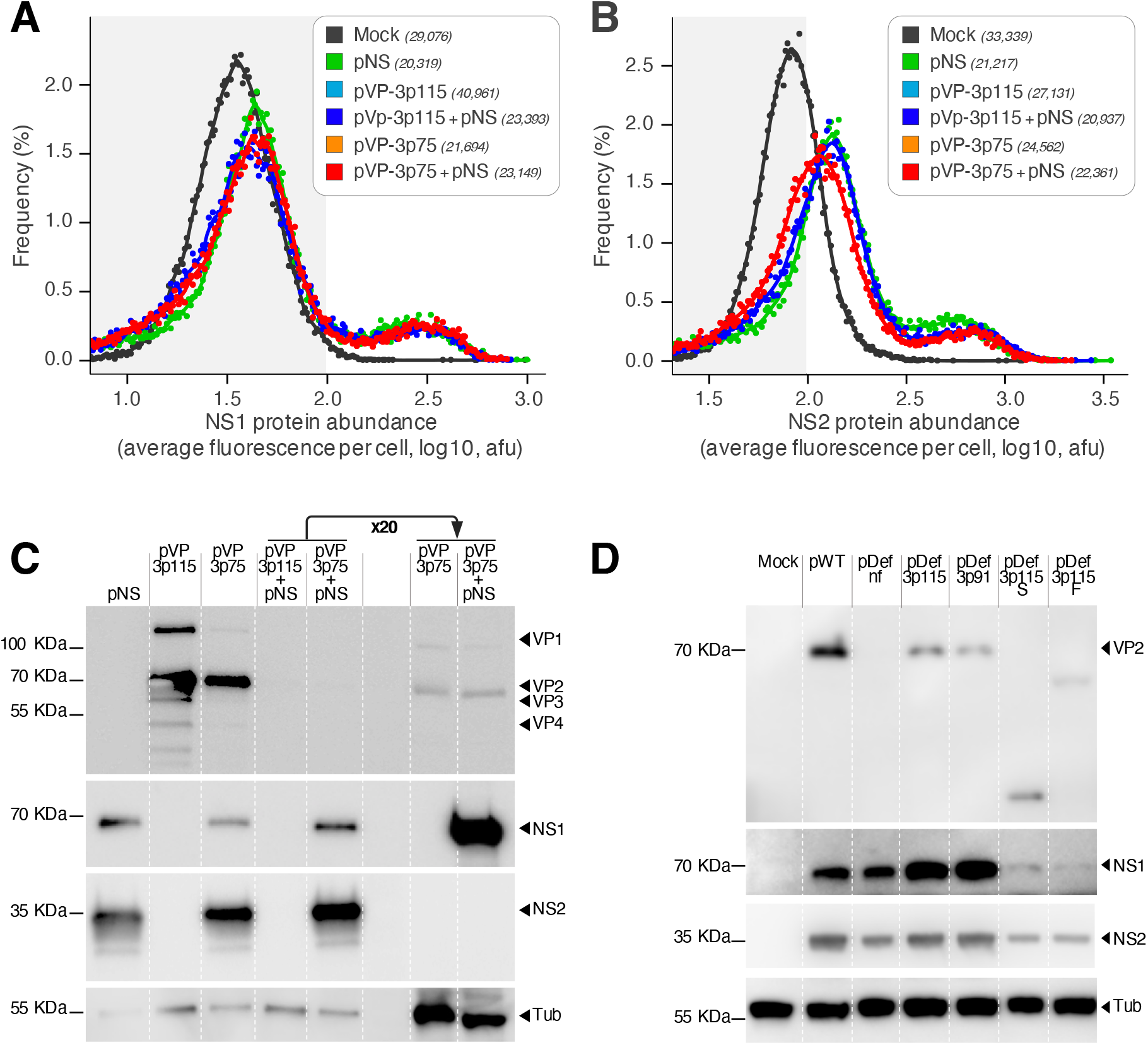
Alteration of *vp*’s 3’UTR impacts VP but not NS expression. **AB.** Histograms show distributions of cellular fluorescence intensities, as quantified by immunofluorescence microscopy for NS1 (**A**) and NS2 (**B**) proteins 3 days after transfection of Ld652Y cells by different combination of *vp* and *ns* blocks, as shown. Numbers indicate the number of analyzed cells. Data points represent cell frequencies over each percentile of the fluorescence range. For visual clarity, points are underlaid with loess regression smoothers. The 99^th^ percentile of control cells (grey background) is used as a threshold to quantify positive fluorescence signals. Barplots showing the sum of fluorescence intensities for cells above threshold are showed in Figure 5C. **CD.** Western blots marked by primary antibodies against VP (top), NS1 (middle-top), NS2 (middle-bottom) and α-Tubulin (bottom), as shown. Protein species are indicated by arrows on the right of the panel. Apparent molecular weights are indicated on the right. Samples are denatured total protein extracted from Ld652Y cells 3 days after transfection with different plasmids, as indicated on top of each lane. Unless stated otherwise, all samples were diluted to the same extent prior to loading. Images were acquired using a chemiluminescence imager, and have been cropped to the signal of interest and individually adjusted for brightness and contrast. Panel **C** shows results for the complementation assay between non-replicable genome blocks (panel **AB** and **Figure 5BC)**. In the last two lanes, samples were diluted 20-fold less for comparison purposes. Panel **D** show results for potentially replicable genomes, including the WT and non-propagatable clones.

**Supplementary Figure 5.**
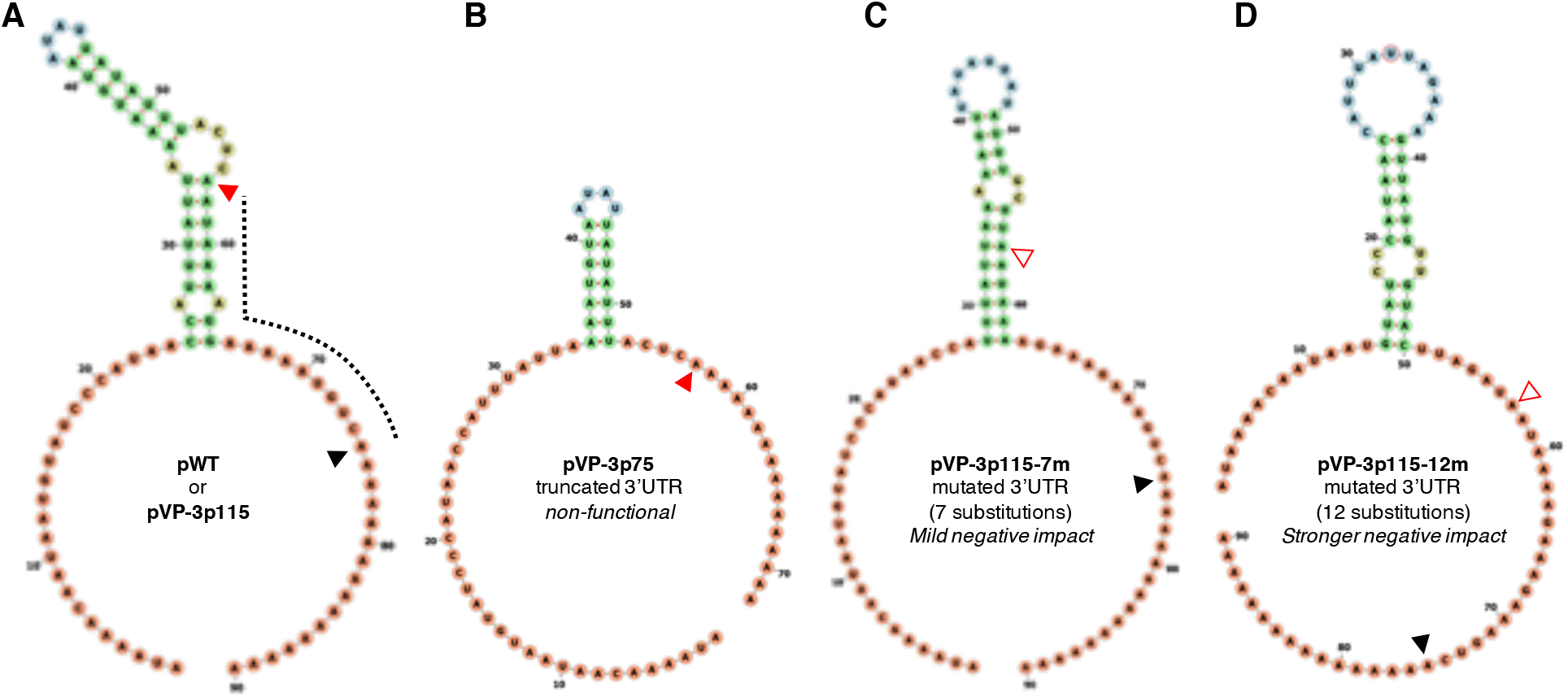
Potential structural binding site for regulatory protein in the 3’UTR of *vp*’s transcript. **A.** Secondary structure prediction in the 3’UTR of *vp*’s native transcript. The 75 nts long 3’UTR was appended with a 15 nts polyA tail and folded. Bases are colored according to structural status. The sequence absents from early-terminated transcript that are non-functional for replication is highlighted with a dotted line: 8 nts are involved in the secondary structure. **B.** Secondary structure prediction in the 3’UTR of *vp*’s truncated transcript (from *e.g.* p118, Figure 4). The lower stem of the structure can no longer be formed. **CD.** Secondary structure prediction from *vp*’s 3’UTR mutants originally constructed to test antisense regulation. The 3’UTR is assumed to have WT length because the cryptic polyA site is mutated, though this was not verified experimentally. Mutations (7 and 12 in pVP-3p115-7m and pVP-3p115-12m, respectively) results in increasingly altered secondary structures in both strength and position, which may explain the increasingly negative impact on replication associated with these variants (Figure 3). Secondary structures were predicted using the Vienna packages RNAfold webserver and rendered with FORNA (47). Black filled triangles mark the WT polyA, red filled triangle the cryptic polyA site and hollow red triangles mutated cryptic polyA site.

**Supplementary Table 1.** Complete plasmids list.

| Full ID | Full name | Description | Reference |
| --- | --- | --- | --- |
| pGCDV1 | pBRJH | Functional WT JcDV clone | 11 |
| pGCDV3 | pITR-A3-GFP | *B. mori*'s A3 actin promoter driving eGFP followed by SV40 polyA signals. Flanked by JcDV's ITR. | 11 |
| pGCDV4 | pNeo | Contains neomycin resistance gene (neo) | Amersham |
| pGCDV10 | pSB1C3::T7polymerase | Cloning backbone pSB1C3 with T7 polymerase insertion | iGEM DNA distribution (2014) |
| pGCDV14 | pBRJH-CmR | WT JcDV clone derived from the original pBRJH clone | This work |
| pGCDV17 | p15aCmR-eGFP-A3-Neo | Promoterless eGFP followed by SV40 polyadenylation signals and *B. mori*'s A3 promoter driving *neo* gene. | This work |
| pGCDV18 | pBbA2c-RFP | Minimal cloning backbone with p15a ori and chloramphenicol resistance | 13 |
| pGCDV24 | mCerulean3-mVenus-FRET-10 | A plasmid comprising mCerulean and mVenus coding sequences | addgene #58180 |
| pGCDV25 | pIZ-Flag6His-Siwi | pIZ backbone with Siwi gene insertion | 15 (Addgene #50560) |
| pGCDV36 | p15aCmR-mVenus-A3-Neo | Promoterless mVenus followed by SV40 polyadenylation signals and *B. mori*'s A3 promoter driving *neo* gene. Flanked by BsaI sites. | This work |
| pGCDV39 | pUC57-Kan-synITR_oxford | Extended ITR from oxford JcDV genome sequence synthesized by Genewiz and cloned in pUC57-Kan. Flanked by BsaI sites | 14 |
| pGCDV40 | p15aCmR-vp4 | JcDV's vp4. Flanked by BsmBI sites | This work |
| pGCDV42 | p15aCmR-P9-vp123-2xBsmBI | JcDV's P9 and vp sequence excluding vp4 which is replaced by two BsmBI accepting sites. Flanked by BsaI sites. | This work |
| pGCDV45 | pIZ-ns-rmBsaI | pIZ backbone with ns gene block under POPEI-2 baculovirus promoter | This work |
| pGCDV46 | p15aCmR_P93 | JcDV's P93 promoter. Flanked by BsaI sites. | This work |
| pGCDV47 | p15aCmR_synITR_oxford | Extended ITR from oxford JcDV genome sequence subcloned in p15a-CmR backbone. Flanked by BsaI sites. | This work |
| pGCDV49 | pIZ-ns-BsaI_Lp | pIZ backbone with ns gene block under POPEI-2 baculovirus promoter and BsaI landing pad | This work |
| pGCDV55 | pFAB217 | A bacterial expression plasmid | 18 |
| pGCDV57 | NoPropJcDV-venus-no_vp4 | Original design of JcDV's non-propagatable clone. No vp4 sequence but BsmBI landing pad. | This work |
| pGCDV58 | NoPropJcDV-venus-vp4-16_tail | Original (non-functional) design of JcDV's non-propagatable clone with 16 nts after vp's stop codon | This work |
| pGCDV73 | NoPropJcDV-venus-vp4-Δns | A derivative of p58 in which the *ns* gene block is replaced by a 500+ nts DNA spacer | This work |
| pGCDV90 | pJDvp | JcDV's P93 and *ns* followed by 484 nts after *ns1*'s stop codon | 17 |
| pGCDV91 | pJDns | JcDV's P9 and *vp* followed by 115 nts after *vp*'s stop codon | 17 |
| pGCDV92 | pJA | pBRJH derivative with truncated ITRs (no hairpin, no replication) | 16 |
| pGCDV93 | pJDns_49_tail | JcDV's P9 and *vp* followed by 49 nts after *vp*'s stop codon | This work |
| pGCDV103 | NoPropJcDV-venus-vp4-115_tail | Amended (functional) design of JcDV's non-propagatable clone with 115 nts after *vp*'s stop codon | This work |
| pGCDV104 | NoPropJcDV-venus-vp4-91_tail | Amended (functional) design of JcDV's non-propagatable clone with 91 nts after *vp*'s stop codon | This work |
| pGCDV105 | p15aCmR-vp4-115_tail | JcDv's vp4 followed by 115 nts after stop codons. Flanked by BsmBI sites | This work |
| pGCDV106 | p15aCmR-vp4-91_tail | JcDv's vp4 followed by 91 nts after stop codons. Flanked by BsmBI sites | This work |
| pGCDV116 | JcDV_Replication_re<br>porter | JcDV's extended (oxford) ITRs flanking a DNA sequence of the same length as the WT genome. Comprises eGFP driven by *B. Mori*'s A3 promoter and followed by SV40 polyadenylation signal | This work |
| pGCDV118 | pJDns-75_tail | JcDv's P9 and *vp* followed by 75 nts after *vp*'s stop codon (stops just after polyA site) | This work |
| pGCDV144 | pJDns-7m_ns_recod | JcDv's P9 and *vp* followed by 115 nts after *vp*'s stop codon with 7 substitutions in the ns1 overlapping region | This work |
| pGCDV145 | pJDns-12m_ns_recod | JcDv's P9 and *vp* followed by 115 nts after *vp*'s stop codon with 12 substitutions in the ns1 overlapping region | This work |
| pGCDV146 | pJDvp-7m_ns_recod | JcDv's P93 and ns followed by 484 nts after ns1's stop codon. 7 silent substitutions in the end of ns1 | This work |
| pGCDV147 | pJDvp-12m_ns_recod | JcDv's P93 and ns followed by 484 nts after ns1's stop codon. 12 silent substitutions in the end of ns1 | This work |
| pGCDV252 | p15aCmR-vp4_168tag-115_tail | JcDv's vp4 followed by 115 nts after stop codons and tat542tag substitution in *vp* (stop codon). Flanked by BsmBI sites | This work |
| pGCDV253 | p15aCmR-vp4_314aa--115_tail | JcDv's vp4 followed by 115 nts after stop codons and aat687aa-deletion in *vp* (frameshift). Flanked by BsmBI sites | This work |
| pGCDV254 | DefectJcDV-venus-vp4_168tag-115_tail | Amended design of JcDV's non-propagatable clone with 115 nts after *vp*'s stop codon and tat542tag substitution in *vp* (stop codon) | This work |
| pGCDV255 | DefectJcDV-venus-vp4_314aa--115_tail | Amended design of JcDV's non-propagatable clone with 115 nts after *vp*'s stop codon and aat687aa- deletion in *vp* (frameshift) | This work |

**Supplementary Table 2.**
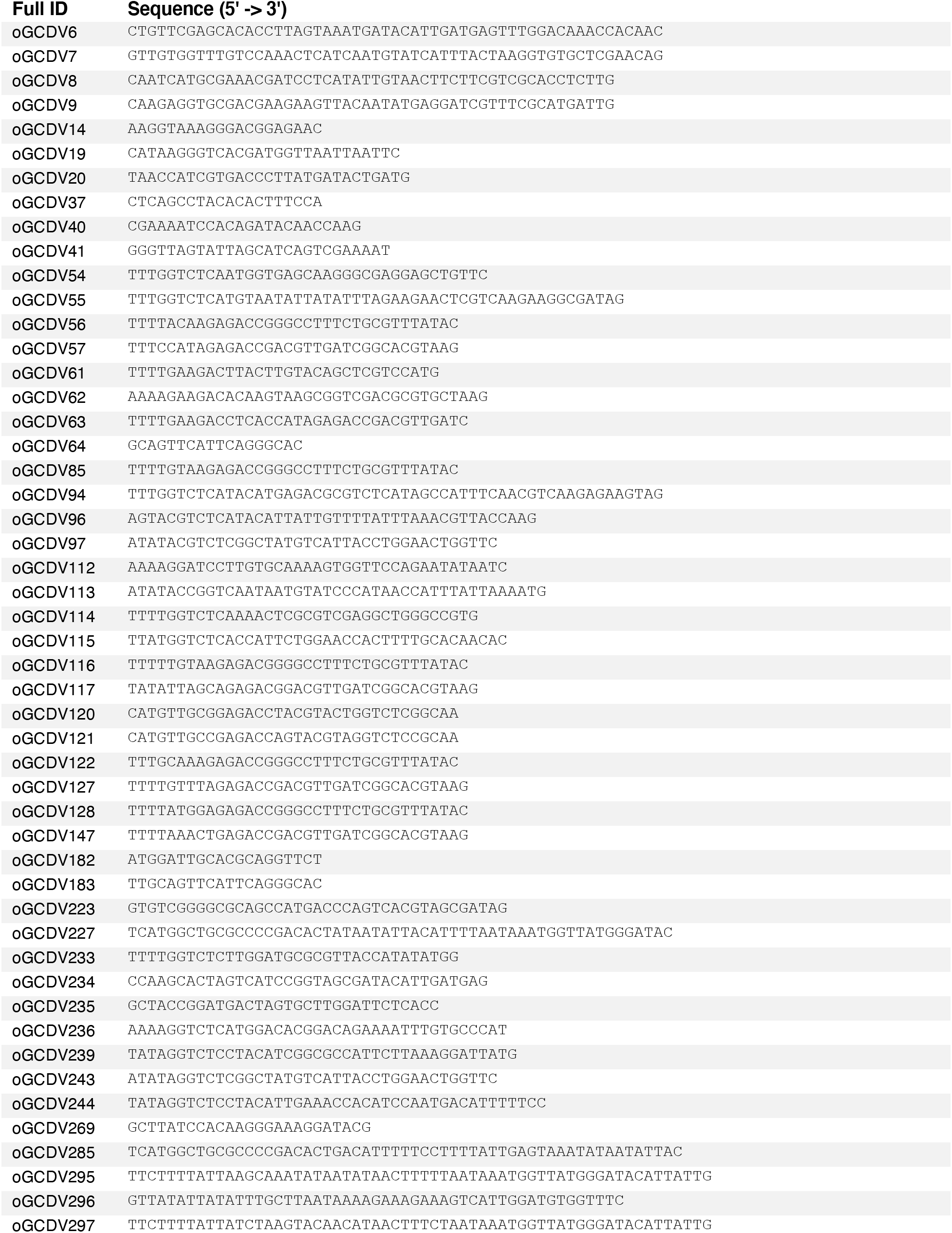
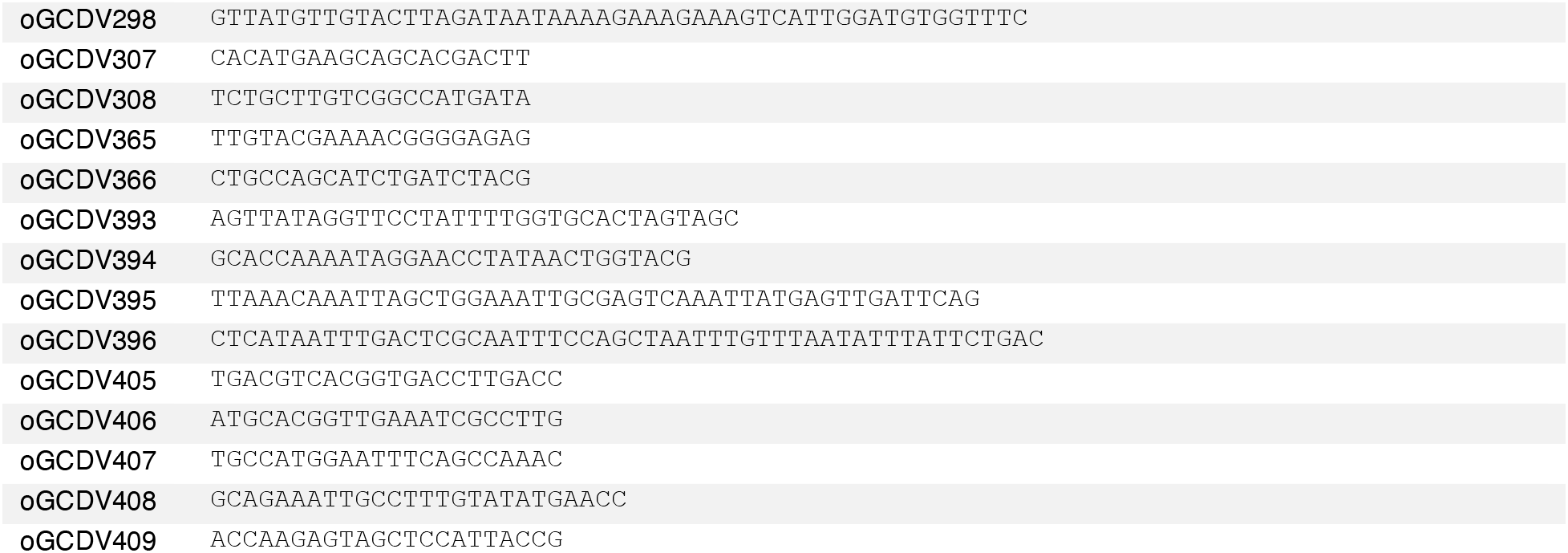
Oligonucleotides.

## References

1. Koonin EV, Dolja VV, Krupovic M. 2015. Origins and evolution of viruses of eukaryotes: The ultimate modularity. Virology 479-480:2–25.

2. Fakhiri J, Schneider MA, Puschhof J, Stanifer M, Schildgen V, Holderbach S, Voss Y, Andari El J, Schildgen O, Boulant S, Meister M, Clevers H, Yan Z, Qiu J, Grimm D. 2019. Novel Chimeric Gene Therapy Vectors Based on Adeno-Associated Virus and Four Different Mammalian Bocaviruses. Molecular Therapy - Methods & Clinical Development 12:202–222.

3. Kerr J, Cotmore S, Bloom ME, Linden RM, Parrish CR. 2006. Parvoviruses. Hodder Arnold.

4. Matthews G. 2018. The spread of fall armyworm (FAW) Spodotera frugiperda. Outlooks on Pest Management 29:213–214.

5. Bergoin M, Tijssen P. 1998. Biological and molecular properties of densoviruses and their use in protein expression and biological control 141–169.

6. Carlson J, Suchman E, Buchatsky L. 2006. Densoviruses for Control and Genetic Manipulation of Mosquitoes. Adv Virus Res 68:361–392.

7. Tijssen P. 1999. Molecular and structural basis of the evolution of parvovirus tropism. Acta Vet Hung 47:379–394.

8. Allison AB, Organtini LJ, Zhang S, Hafenstein SL, Holmes EC, Parrish CR, Ross SR. 2016. Single Mutations in the VP2 300 Loop Region of the Three-Fold Spike of the Carnivore Parvovirus Capsid Can Determine Host Range. J virol, 3rd ed. 90:753–767.

9. Multeau C, Froissart R, Perrin A, Castelli I, Casartelli M, Ogliastro M. 2012. Four amino acids of an insect densovirus capsid determine midgut tropism and virulence. J virol 86:5937–5941.

10. Marsic D, Govindasamy L, Currlin S, Markusic DM, Tseng Y-S, Herzog RW, Agbandje-McKenna M, Zolotukhin S. 2014. Vector Design Tour de Force: Integrating Combinatorial and Rational Approaches to Derive Novel Adeno-associated Virus Variants. Mol Ther 22:1900–1909.

11. Nonnenmacher M, van Bakel H, Hajjar RJ, Weber T. 2015. High Capsid–Genome Correlation Facilitates Creation of AAV Libraries for Directed Evolution. Mol Ther 23:675–682.

12. Ogden PJ, Kelsic ED, Sinai S, Church GM. 2019. Comprehensive AAV capsid fitness landscape reveals a viral gene and enables machine-guided design. Science 366:1139–1143.

13. Bossin H. 1998. Développement de vecteurs d“expresssion stable dérivés du densovirus JcDNV: application à l”expression constitutive de protéines hétérologues en lignées cellulaires de lépidoptères et comme marqueur au cours du développement chez la drosophile. Montpellier 2.

14. Engler C, Kandzia R, Marillonnet S. 2008. A one pot, one step, precision cloning method with high throughput capability. PLoS ONE 3:e3647.

15. Lee TS, Krupa RA, Zhang F, Hajimorad M, Holtz WJ, Prasad N, Lee SK, Keasling JD. 2011. BglBrick vectors and datasheets: A synthetic biology platform for gene expression. J Biol Eng 5:12.

16. Pham HT, Huynh OTH, Jousset FX, Bergoin M, Tijssen P. 2013. Junonia coenia Densovirus (JcDNV) Genome Structure. Genome Announcements 1:e00591-13–e00591-13.

17. Kawaoka S, Hayashi N, Suzuki Y, Abe H, Sugano S, Tomari Y, Shimada T, Katsuma S. 2009. The Bombyx ovary-derived cell line endogenously expresses PIWI/PIWI-interacting RNA complexes. RNA 15:1258–1264.

18. Li Y, Jousset F, Giraud C, Rolling F, Bergoin M. 1996. A titration procedure of the Junonia cœia densovirus and quantitation of transfection by its cloned genomic DNA in four lepidopteran cell lines. Journal of virological….

19. Salasc F, Mutuel D, Debaisieux S, Perrin A, Dupressoir T, Grenet ASG, Ogliastro M. 2016. Role of the phosphatidylinositol-3-kinase/Akt/target of rapamycin pathway during ambidensovirus infection of insect cells. Journal of General Virology 97:233–245.

20. Mutalik VK, Guimaraes JC, Cambray G, Mai Q-A, Christoffersen MJ, Martin L, Yu A, Lam C, Rodriguez C, Bennett G, Keasling JD, Endy D, Arkin AP. 2013. Quantitative estimation of activity and quality for collections of functional genetic elements. Nat Methods 10:347–353.

21. Cambray G, Guimaraes JC, Arkin AP. 2018. Evaluation of 244,000 synthetic sequences reveals design principles to optimize translation in *Escherichia coli*. Nat Biotechnol 54:198.

22. Goodwin RH, Tompkins GJ, McCawley P. 1978. Gypsy moth cell lines divergent in viral susceptibility. I. Culture and identification. In Vitro 14:485–494.

23. Wang Y, Abd-Alla AMM, Bossin H, Li Y, Bergoin M. 2013. Analysis of the transcription strategy of the Junonia coenia densovirus (JcDNV) genome. Virus Res 174:101–107.

24. Cotmore SF, Tattersall P. 2014. Parvoviruses: Small Does Not Mean Simple. Annu Rev Virol 1:517–537.

25. Brandenburger A, Coessens E, Bakkouri KE, Velu T. 1999. Influence of Sequence and Size of DNA on Packaging Efficiency of Parvovirus MVM-Based Vectors. Human Gene Therapy 10:1229–1238.

26. Bleckmann M, Schürig M, Chen F-F, Yen Z-Z, Lindemann N, Meyer S, Spehr J, van den Heuvel J. 2016. Identification of Essential Genetic Baculoviral Elements for Recombinant Protein Expression by Transactivation in Sf21 Insect Cells. PLoS ONE 11:e0149424.

27. Proudfoot NJ. 2011. Ending the message: poly(A) signals then and now. Genes & Development 25:1770–1782.

28. Jacobson A, Peltz SW. 1996. Interrelationships of the Pathways of mRNA Decay and Translation in Eukaryotic Cells. Annu Rev Biochem 65:693–739.

29. Giudice G, Sánchez-Cabo F, Torroja C, Lara-Pezzi E. 2016. ATtRACT-a database of RNA-binding proteins and associated motifs. Database (Oxford) 2016:baw035.

30. Szostak E, Gebauer F. 2013. Translational control by 3’-UTR-binding proteins. Briefings in Functional Genomics 12:58–65.

31. Mazumder B, Poddar D, Basu A, Kour R, Verbovetskaya V, Barik S, García-Sastre A. 2014. Extraribosomal L13a Is a Specific Innate Immune Factor for Antiviral Defense. J virol 88:9100–9110.

32. Rhode SL, Richard SM. 1987. Characterization of the trans-activation-responsive element of the parvovirus H-1 P38 promoter. J virol 61:2807–2815.

33. Li X, Rhode SL III. 1993. The Parvovirus H-1 NS2 Protein Affects Viral Gene Expression through Sequences in the 3′ Untranslated Region. Virology 194:10–19.

34. Christensen J, Cotmore SF, Tattersall P. 1995. Minute virus of mice transcriptional activator protein NS1 binds directly to the transactivation region of the viral P38 promoter in a strictly ATP-dependent manner. J virol 69:5422–5430.

35. Ward TW, Kimmick MW, Afanasiev BN, Carlson JO. 2001. Characterization of the Structural Gene Promoter of Aedes aegypti Densovirus. J virol 75:1325–1331.

36. Raab U, Beckenlehner K, Lowin T, Niller H-H, Doyle S, Modrow S. 2002. NS1 Protein of Parvovirus B19 Interacts Directly with DNA Sequences of the p6 Promoter and with the Cellular Transcription Factors Sp1/Sp3. Virology 293:86–93.

37. Ding C, Urabe M, Bergoin M, Kotin RM. 2002. Biochemical Characterization of Junonia coenia Densovirus Nonstructural Protein NS-1. J virol 76:338–345.

38. Abd-Alla A, Jousset F-X, Li Y, Fédière G, Cousserans F, Bergoin M. 2004. NS-3 protein of the Junonia coenia densovirus is essential for viral DNA replication in an Ld 652 cell line and Spodoptera littoralis larvae. J virol 78:790–797.

39. Powell SK, Samulski RJ, McCown TJ. 2020. AAV Capsid-Promoter Interactions Determine CNS Cell-Selective Gene Expression In Vivo. Mol Ther 28:1373–1380.

40. Mao Y, Su J, Wang J, Zhang X, Hou Q, Bian D, Liu W. 2016. Roles of three amino acids of capsid proteins in mink enteritis parvovirus replication. Virus Res 222:24–28.

41. Fernandes S, Boisvert M, Tijssen P. 2011. Genetic Elements in the VP Region of Porcine Parvovirus Are Critical to Replication Efficiency in Cell Culture. J virol 85:3025–3029.

42. Ward P, Clément N, Linden RM. 2007. cis Effects in Adeno-Associated Virus Type 2 Replication. J virol 81:9976–9989.

43. Saxena P, Lomonossoff GP. 2014. Virus Infection Cycle Events Coupled to RNA Replication. Annu Rev Phytopathol 52:197–212.

44. Wu Z, Yang H, Colosi P. 2009. Effect of Genome Size on AAV Vector Packaging. Mol Ther 18:80–86.

45. Cui J, Schlub TE, Holmes EC. 2014. An Allometric Relationship between the Genome Length and Virion Volume of Viruses. J virol 88:6403–6410.

46. Lucía-Sanz A, Manrubia S. 2017. Multipartite viruses: adaptive trick or evolutionary treat? npj Syst Biol Appl 3:1–11.

47. Kerpedjiev P, Hammer S, Hofacker IL. 2015. Forna (force-directed RNA): Simple and effective online RNA secondary structure diagrams. Bioinformatics 31:3377–3379.

